# Single-nucleus transcriptome analysis reveals cell type-specific molecular signatures across reward circuitry in the human brain

**DOI:** 10.1101/2020.10.07.329839

**Authors:** Matthew N. Tran, Kristen R. Maynard, Abby Spangler, Leonardo Collado-Torres, Vijay Sadashivaiah, Madhavi Tippani, Brianna K. Barry, Dana B. Hancock, Stephanie C. Hicks, Joel E. Kleinman, Thomas M. Hyde, Keri Martinowich, Andrew E. Jaffe

**Author notes:** Equal contributions (Matthew N. Tran, Kristen R. Maynard). Co-corresponding authors; / @martinowk, / @andrewejaffe.

## Abstract

Single cell/nucleus technologies are powerful tools to study cell type-specific expression in the human brain, but most large-scale efforts have focused on characterizing cortical brain regions and their constituent cell types. However, additional brain regions - particularly those embedded in basal ganglia and limbic circuits - play important roles in neuropsychiatric disorders and addiction, suggesting a critical need to better understand their molecular characteristics. We therefore created a single-nucleus RNA-sequencing (snRNA-seq) resource across five human brain regions (hippocampus, HPC; dorsolateral prefrontal cortex, DLPFC; subgenual anterior cingulate cortex, sACC; nucleus accumbens, NAc; and amygdala, AMY), with emphasis on the NAc and AMY, given their involvement in reward signaling and emotional processing. We identified distinct and potentially novel neuronal subpopulations, which we validated by smFISH for various subclasses of NAc interneurons and medium spiny neurons (MSNs). We additionally benchmarked these datasets against published datasets for corresponding regions in rodent models to define cross-species convergence and divergence across analogous cell subclasses. We characterized the transcriptomic architecture of regionally-defined neuronal subpopulations, which revealed strong patterns of similarities in specific neuronal subclasses across the five profiled regions. Finally, we measured genetic associations between risk for psychiatric disease and substance use behaviors with each of the regionally-defined cell types. This analysis further supported NAc and AMY involvement in risk for psychiatric illness by implicating specific neuronal subpopulations, and highlighted potential involvement of an MSN population associated with stress signaling in genetic risk for substance use.

## Introduction

Recent advances in single-cell and single-nucleus RNA-sequencing (scRNA-seq/snRNA-seq) technologies have facilitated the molecular characterization of diverse cell types in the postmortem human brain during development (Darmanis et al., 2015; Li et al., 2018a; Zhong et al., 2018, 2020), and have been used to assess cell type-specific gene expression differences in the context of several brain disorders, including Alzheimer’s disease, autism spectrum disorder, multiple sclerosis, and major depressive disorder (Mathys et al., 2019; Nagy et al., 2020; Schirmer et al., 2019; Velmeshev et al., 2019). Identification of cell type-specific gene expression signatures has contributed to the understanding of the relationship between molecular identity and cell function as it relates to brain health, neurological disease, and genetic risk for neuropsychiatric disorders, such as schizophrenia (Skene et al., 2018).

While substantial advancements have been made in understanding cell type heterogeneity within regions and across the human brain, the majority of snRNA-seq reports are limited to a small number of brain areas. These primarily include the hippocampus (HPC) (Franjic et al., 2020; Habib et al., 2017) and several heavily studied sub-regions of the cortex (Lake et al., 2016), including the dorsolateral prefrontal cortex (DLPFC) (Li et al., 2018a; Nagy et al., 2020), medial temporal cortex (Darmanis et al., 2015; Hodge et al., 2019), entorhinal cortex (Grubman et al., 2019), and anterior cingulate cortex (Velmeshev et al., 2019). Molecular profiling of less studied cortical subregions including the subgenual anterior cingulate cortex (sACC), as well as striatal and limbic brain regions, including the nucleus accumbens (NAc) and the amygdala (AMY), is lacking in the human brain. The sACC, NAc, and AMY are interconnected within well-established circuit loops that mediate important behavioral and neurobiological functions, including signaling for reward and motivation as well as processing emotional valence, particularly for fearful and threatening stimuli (Haber and Knutson, 2010; Janak and Tye, 2015; Russo and Nestler, 2013). Importantly, the cellular composition of individual neuronal subtypes in these regions substantially differs from previously well-profiled cortical and hippocampal regions (Saunders et al., 2018; Zeisel et al., 2018).

For example, the NAc contains dopaminoceptive populations of GABAergic medium spiny neurons (MSNs) - the principal projecting cell type comprising up to 95% of neurons in rodent - that harbor unique physiological and cellular properties (Gerfen et al., 1990; Kawaguchi, 1997; Kronman et al., 2019; Russo and Nestler, 2013). Early functional characterization of MSNs revealed two distinct classes of MSNs based on expression of D1 versus the D2 dopamine receptors (D1-MSNs and D2-MSNs, respectively) (Lobo, 2009; Lobo et al., 2006). However, recent sc/sn-RNAseq studies in the rodent striatum, and in the NAc specifically, revealed more complex transcriptional diversity within broader D1 and D2-MSN subclasses than was previously appreciated (Gokce et al., 2016; Saunders et al., 2018; Stanley et al., 2020; Zeisel et al., 2018). Moreover, subpopulations of MSNs are differentially recruited in response to cocaine exposure, and mediate divergent functional effects on behavioral responses to drugs of abuse (Savell et al., 2020). Similarly, single-cell profiling studies in the rodent AMY identified specialized populations of *Cck*-expressing neurons that are preferentially activated by behavioral experience, including exposure to acute stress (Wu et al., 2017). However, whether and to what extent this transcriptional diversity is conserved in these areas of the human NAc and amygdala has not yet been fully explored. Given evidence for the functional importance of unique cell types in these areas of the rodent brain, profiling these regions in human by snRNA-seq may identify analogous cell populations, which can then be analyzed in the context of neurobiological dysfunction in human brain disorders.

Further, while unique transcriptomic profiles have been identified for specialized cell types that are localized to specific brain regions (e.g. granule cells of the dentate gyrus; MSNs of the striatum), it remains unclear to what extent broad populations of “common” cell types, such as glutamatergic excitatory neurons, are transcriptionally similar both within and across different brain regions. Given that many snRNA-seq studies have used different nuclei isolation protocols (sucrose vs iodixanol gradients), cell enrichment techniques (NeuN or DAPI sorting), library/sequencing technologies (10x Genomics Chromium, SMARTseq, Fluidigm, DroNc-seq), and data analysis workflows (dimensionality reduction techniques, handling batch effects, cluster annotation), comparing cell type-specific signatures across studies and between brain regions has been computationally and methodologically challenging. While efforts to generate a human cell type atlas at the single-cell level are underway (Han et al., 2020), the landscape of specialized molecular cell types across the human brain remains largely unexplored.

Here we defined the molecular taxonomy of distinct cell types in subcortical regions (NAc and AMY), which act as key nodes within circuits that mediate critical brain and behavioral functions including reward signaling and emotional processing. We also validated molecular profiles for previously identified cell types in the HPC and DLPFC, and identify similar cell types in the sACC, an additional cortical region central to limbic system function that has been implicated in affective disorders. Furthermore, we evaluate cross-species conservation of NAc and AMY cell types between human and rodent, specifically focusing on comparisons of MSN sub-populations identified as playing key roles in reward-processing and addiction. Finally, by integrating genetic studies for substance use and neuropsychiatric disorders, we show differential cell type expression of genes associated with schizophrenia, autism spectrum disorder, major depressive disorder, bipolar disorder, posttraumatic stress disorder, and features of addiction, highlighting the clinical relevance of understanding cell type- and regionspecific expression in the human brain.

## Results

We profiled 5 brain regions (DLPFC, HPC, sACC, NAc, and AMY) across up to 5 neurotypical, adult male subjects of European Ancestry using 10x Genomics Chromium technology. To minimize potential batch effects, regions/donors were split across Chromium runs, for a total of 14 samples (sample/demographic information found in **Table S1**). Nuclear preparations were generated and purified by flow cytometry using DAPI staining (and NeuN selection for a subset of samples) to obtain nuclei from all cell types in a brain region. After sequencing, data processing and QC (Methods; **Table S2**), we analyzed a total of 42,308 high-quality nuclei across these 14 samples, which were then analyzed in respective region-specific, in addition to pan-brain, analyses.

### Identification of refined medium spiny neuron subpopulations in human NAc

To evaluate the transcriptional landscape of MSNs and other cell populations in the human NAc, we analyzed 13,148 total nuclei from 5 donors, including 4,465 DAPI-sorted nuclei and 8,683 NeuN-sorted nuclei, which allowed for enrichment of MSNs. We performed data-driven clustering to generate 14 cell subclusters across six broad cell types, including GABAergic inhibitory interneurons, MSNs, oligodendrocytes, oligodendrocyte precursor cells, microglia, and astrocytes (**Figure 1A**). Of the 6 distinct neuronal clusters expressing established D1- and D2-MSNs markers (**Figure 1B**), including *PPP1R1B* (encoding DARPP-32), four of these MSN subclusters were enriched for *DRD1* (D1.1, D1.2, D1.3, D1.4) and two were enriched for *DRD2* (D2.1 and D2.2). These MSN subclusters collectively made up 94% and 95% of neuronal nuclei from the two neuron-enriched samples (**Table S3**), lending human evidence that the vast majority of nuclei in this region of the striatum are composed of MSNs, as in rodent (Kawaguchi, 1997). Clusters D1.4 and D2.2 represented the largest D1-MSN (77%) and D2-MSN (92%) subclasses, respectively. As expected, MSN subclusters showed differential enrichment of several neuropeptides, including proenkephalin (*PENK*), tachykinin 1 (*TAC1*), and prodynorphin (*PDYN*) (**Figure S1**; top markers listed in **Table S4**) (Lobo, 2009; Lobo et al., 2006; Savell et al., 2020). Surprisingly, the classical D1-MSN marker, *TAC1*, was enriched in D2.1 MSNs, but largely absent in D1.1 and D1.2 MSNs (**Figure 1B**). Similarly, the classical D2-MSN marker *PENK* was enriched in the large population of D2.2 MSNs, but depleted in the smaller population of D2.1 MSNs (**Figure S1**). Expression of these neuropeptides in D1 and D2 MSN subclasses was confirmed using single molecule fluorescent in situ hybridization (smFISH) with 4-plex RNAscope technology (Maynard et al., 2020b), and showed that the majority of D2-expressing cells in the NAc belonged to the D2.2 class (86.8%, **Figure S1**).

**Figure 1:**
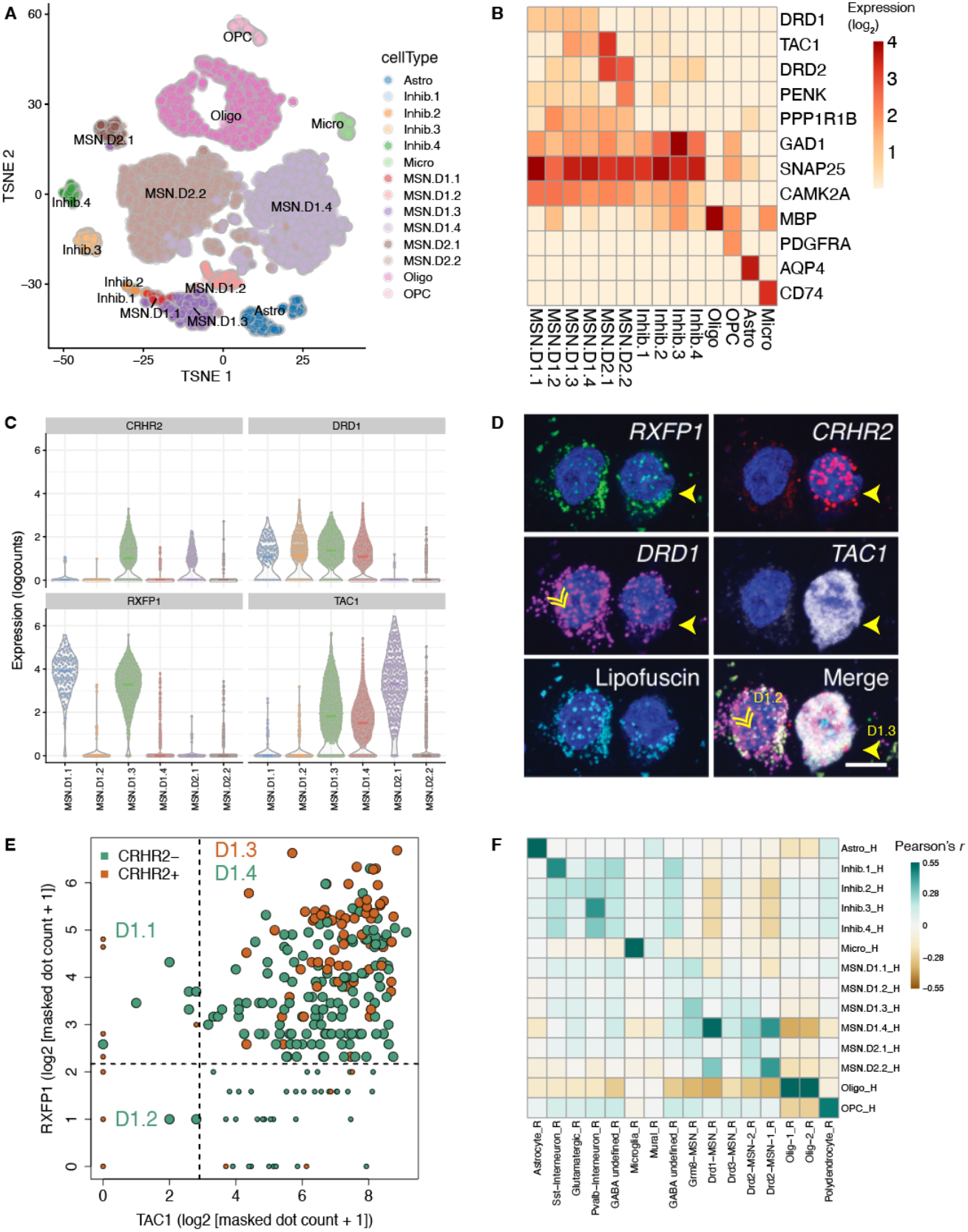
Distinct subpopulations of D1- and D2-expressing MSNs in human NAc. **(A)** tSNE plot of 13,148 nuclei (n=5 donors) across 14 clusters, including 4 clusters of D1 MSNs and 2 clusters of D2 MSNs. (**B**) Heatmap depicting log_2_ expression of known marker genes in each cluster. **(C)** Violin plots for 4 genes differentially expressed (log_2_-normalized counts) in specific D1 subpopulations (*CRHR2, DRD1, RXFP1, and TAC1*) that were selected for validation using single molecule fluorescent in situ hybridization (smFISH). **(D)** Multiplex smFISH in human NAc depicting D1.2 and D1.3 MSNs. Maximum intensity confocal projections showing expression of DAPI (nuclei), *CRHR2, DRD1, TAC1* and lipofuscin autofluorescence. Merged image without lipofuscin autofluorescence. Scale bar=10 μm. **(E)** Log_2_ expression of respective transcript counts per smFISH region of interest (ROI), post lipofuscin-masking (autofluorescence). Points are colored by *CRHR2* expression and are enlarged where the Euclidean distance = 0 for prediction of MSN subclass for that ROI. **(F)** Heatmap of Pearson correlation values evaluating the relationship between our human-derived NAc statistics (rows) for 14,121 genes and data from (Savell et al., 2020) derived from rat NAc with data-driven clusters provided in their processed data.

Using differential expression analyses, we identified the most preferentially expressed genes in each MSN subcluster and found tens to hundreds of unique markers for D1 and D2-MSN subclasses (at false discovery rate, or FDR, < 1e-6; **Table S4**). Among D1-MSNs, *TAC1*-negative D1.1 MSNs and *TAC1*-positive D1.3 MSNs were both enriched for the GABA_A_ receptor subunit, *GABRQ*, and the relaxin family peptide receptor 1, *RXFP1* (**Figure 1C; Figure S2**). However, only D1.3 MSNs expressed substantial levels of *CRHR2*, encoding corticotropin releasing hormone receptor 2, a protein implicated in mediating the response to stress in the brain (**Figure 1D**). Likewise, *TAC1*-negative D1.2 MSNs could be distinguished from D1.1 and D1.3 MSNs by elevated expression of relaxin family peptide receptor 2, *RXFP2*, and depletion of both *RXFP1* and *GABRQ* (**Figure S2**). Consistent with the identification of a discrete D2-MSN subpopulation expressing *Htr7* in the mouse striatum (Gokce et al., 2016; Stanley et al., 2020), we identified enrichment of *HTR7* in D2.1 (*TAC1*-positive; *PENK*-negative) MSNs, but not D2.2 (*TAC1*-negative; *PENK*-positive) MSNs (**Figure S3**). Similar to D1.3 MSNs, the *HTR7*-positive D2.2 cluster was the only D2-MSN subpopulation expressing *CRHR2*. The existence of these novel D1 and D2 MSN subpopulations was validated by smFISH on NAc brain sections derived from independent postmortem human brain donors (**Figure 1D-E; Figures S1–3**). Several other genes including *CASZ1, GPR6*, and *EBF1* were differentially expressed in unique D1 and/or D2-MSN subpopulations (**Figure S4**). *CASZ1* was highly enriched in the D1.3 and D2.1 subpopulations, *GPR6* in both D2.1 and D2.2 subpopulations, and *EBF1* in the D1.2 subpopulation.

In addition to describing transcriptional diversity in D1 and D2 MSNs, we also identified 4 subclusters of inhibitory interneurons expressing GABAergic marker genes (*GAD1* and *GAD2*), but depleted for MSN marker genes (**Figure 1B; Figure S5**). These clusters contained different transcriptionally-defined classes, including interneurons expressing somatostatin (*STT*; Inhib.1), neuropeptide Y (*NPY*; Inhib 1), prepronociceptin (*PNOC*; Inhib.1), vasoactive intestinal peptide (*VIP*; Inhib.2, albeit a rare subset of these nuclei), and tachykinin 3 (*TAC3*; Inhib.4; **Figure S5** and **Table S4**). While we did not observe robust expression of parvalbumin (*PVALB*) in any cluster, Inhib.3 showed high expression of *KIT*, encoding the protein c-Kit, which is frequently co-expressed in mouse *Pvalb*/PV-positive GABAergic interneurons (Enterría-Morales et al., 2020). smFISH for *PVALB* and other top marker genes for Inhib.3 (*PTHLH, KIT, GAD1*) confirmed that this GABAergic cluster likely represents PV-expressing interneurons (**Figure S5**).

We next evaluated the conservation of NAc cell types across species by comparing our transcriptional profiles with those generated in a previous study of acute exposure to cocaine, which analyzed a total of 15,631 rat NAc nuclei (Savell et al., 2020). Correlation analyses between our NAc subpopulations with those derived from rat NAc revealed that glial populations, including astrocytes, microglia, oligodendrocytes, and oligodendrocyte progenitor cells, were highly conserved (**Figure 1F**). Inhibitory interneuron populations were also well-correlated across species as rat *Sst*-expressing and likely-*Pvalb*-expressing clusters overlapped with human Inhib.1 and Inhib.3 clusters, respectively. We also observed substantial correlation between rat and human D1 and D2-MSNs, especially between rat *Drd1*-expressing MSNs and human D1.4 MSNs. Further, beyond the expected overlap of rat *Drd2*-expressing MSNs in the human D2.2 MSN subcluster, we additionally saw positive correlations across D1 and D2 MSN subtypes, such that rat *Drd2*-expressing MSNs also showed enrichment in our human D1.4 MSN subcluster. This result is not likely fully explained by co-expression of *DRD1* and *DRD2* in the same nucleus because, while we did find that ~9% of all MSNs expressed some degree of both *DRD1* and *DRD2*, these dual-expressing nuclei were mainly restricted to the D1.3 subcluster, as opposed to D1.4 (**Figure 1B**). Additionally, many of the top markers for either the D1.4 or D2.2 subcluster were highly expressed in both MSN clusters (**Figure S4**), suggesting that the majority of canonically dichotomous D1 or D2 MSNs may be more molecularly similar than previously believed. We did not observe strong enrichment for rat *Drd3*- and *Grm8*-expressing MSNs in any of our human MSN subclusters. Likewise, D1.1 and D1.2 human MSNs did not appear to be well represented in rat MSN subtypes (see Discussion). Taken together, while these data suggest strong overall conservation between rat and human NAc cell types, there appear to be transcriptional features that are unique among specialized subpopulations of rodent and human MSNs.

### Atlas of molecularly-defined cell types in amygdala

The amygdala, a medial structure of the temporal lobe is noted for its role in processing emotional valence, particularly for both fear and reward (Janak and Tye, 2015; Wassum and Izquierdo, 2015). Dysfunction in amygdalar signaling is implicated in major depressive disorder, bipolar disorder and posttraumatic stress disorder (PTSD) (Fenster et al., 2018; Garrett and Chang, 2008; Murray et al., 2011). The human amygdala can be subdivided into a number of distinct regions based on histology, immunohistochemical classifications, connectivity, and neural activation patterns as revealed by functional magnetic resonance imaging (fMRI) of the brain (Barger et al., 2012; Schumann and Amaral, 2005; Sorvari et al., 1995; Tyszka and Pauli, 2016; Zhang et al., 2018). Studies in the rodent and non-human primate amygdala have identified different cell compositions across the amygdala, which likely correspond to differential patterns of synaptic connections between cell types across amygdalar subregions, and with extra-amygdalar brain regions (Chareyron et al., 2011). Hence, it is likely that various cell types with unique molecular signatures also exist within the human amygdala, which can be surveyed by snRNA-seq. We therefore analyzed 6,582 nuclei from the greater amygdala of two adult neurotypical donors to create a molecular taxonomy of cell types in this brain region. We identified 12 clusters that corresponded to four broad glial cell types and eight neuronal subclusters (**Figure 2A; Figure S6**). Glial cell populations were present at similar proportions between the donors (52% and 53% Oligo; 15% and 11% Astro; 13% and 10% Micro; 10% and 9% OPC), but we observed a varied distribution of neuronal subclusters between donors (see Discussion; **Table S3**). Despite this, after correcting for donor batch effects, we identified hundreds of genes enriched in each broad glial and neuronal subcluster at FDR < 1e-6 (top markers shown in **Table S4**).

**Figure 2:**
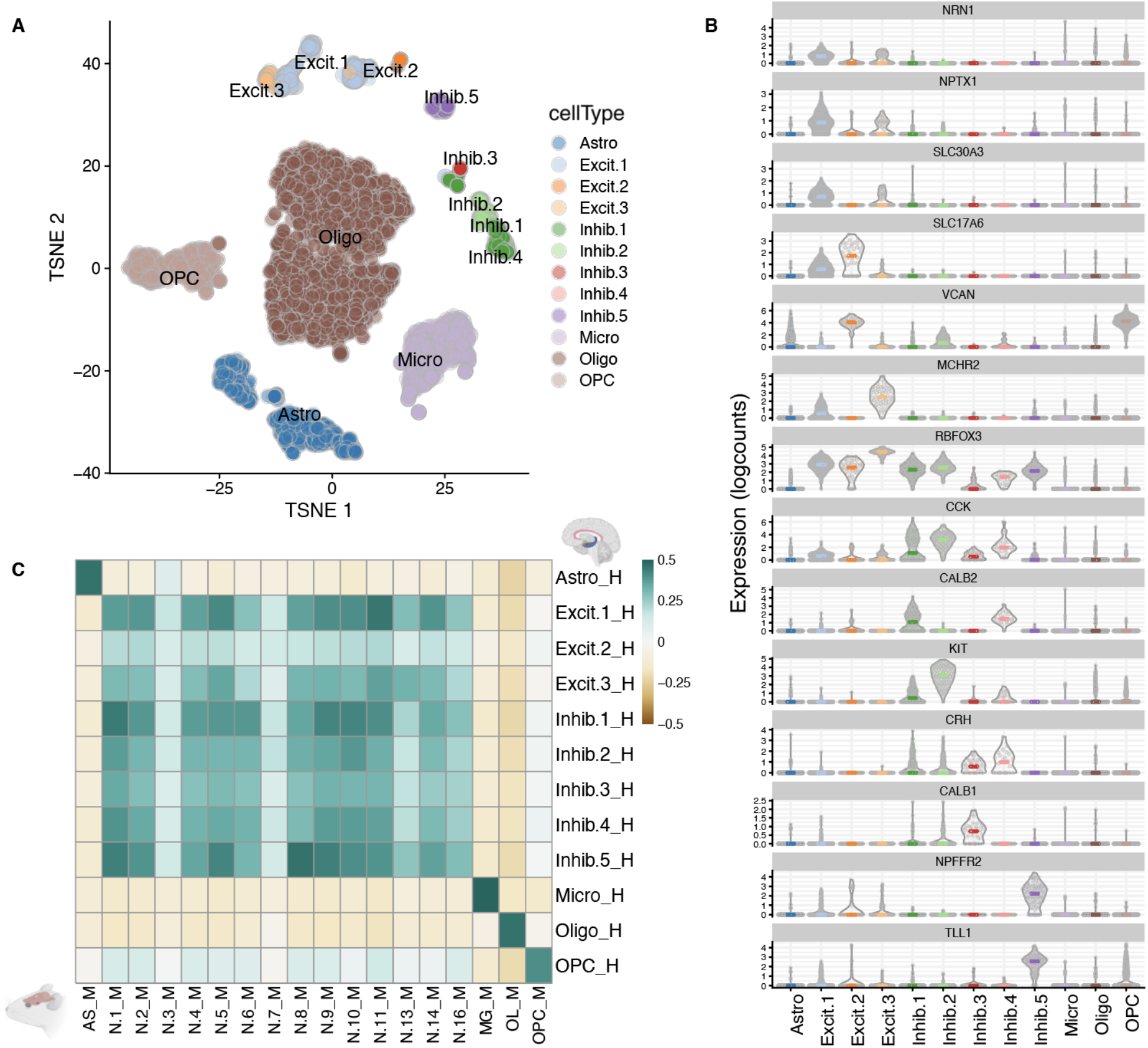
Atlas of molecularly-defined cell types in human AMY. **(A)** tSNE plot of 6,582 nuclei across 12 clusters, including 3 clusters of excitatory neurons and 5 clusters of inhibitory interneurons. **(B)** Expression violin plots for the top 2-3 genes for each of the neuronal subpopulations (log2-normalized counts). **(C)** Heatmap of Pearson correlation values evaluating the relationship between our human-derived amygdala statistics (rows) for 13,525 homologous genes and nuclei from (Chen et al., 2019) derived from mouse amygdala with data-driven clusters provided in their processed data.

Within the eight neuronal subclusters expressing the pan neuronal marker gene *SNAP25*, three clusters were enriched for excitatory neuronal markers (*SLC17A7, SLC17A6*) and five clusters were enriched for inhibitory GABAergic markers (*GAD1, GAD2*; **Figure S6**). The three excitatory subclusters comprised different functional classes of neurons (referred to as Excit.1 to 3), with top markers including *NRN1, NPTX1* and *SLC30A3* (encoding neuritin, neuronal pentraxin 1, and zinc transporter 3, respectively) for Excit.1, and *SLC17A6* and *VCAN* (Versican) for Excit.2 (**Figure 2B)**. *NRN1*, *NPTX1, SLC30A3*, and *VCAN* have all been implicated in modulation of synaptic plasticity and memory (Figueiro-Silva et al., 2015; Horii-Hayashi et al., 2008; Sindreu and Storm, 2011; Yao et al., 2018). The top marker for Excit.3 was *MCHR2* (melanin-concentrating hormone receptor 2), and it was the subpopulation most enriched for *RBFOX3* (NeuN). Compared to excitatory neuron subclusters, we identified a greater diversity of inhibitory GABAergic subpopulations, including *CCK*-containing regular-spiking interneurons (Inhib.1, Inhib.2, Inhib.4) evident by high expression of *CCK* (cholecystokinin; **Figure 2B**). Of these three *CCK*-expressing GABAergic clusters, Inhib.1 and Inhib.4 were also enriched in *CALB2* (calretinin), whereas Inhib.2 showed enrichment for *KIT*. The GABAergic subcluster Inhib.3 was specific for expression of *CALB1* (calbindin), on the other hand, and both Inhib.3 and Inhib.4 were enriched for *CRH* (corticotropin release hormone/factor). CRH is a key regulator of the hypothalamic-pituitary-adrenal (HPA) axis, which is critical for both the acute stress response and adaptation to chronic stress. Finally, *NPFFR2* and *TLL1*, additional genes associated with HPA axis regulation, were selectively expressed in Inhib.5 (Lin et al., 2016; Tamura et al., 2005).

We then compared our subcluster-level transcriptomic profiles to those of a previously published single-cell dataset derived from the mouse medial amygdala (MeA) (Chen et al., 2019) to evaluate conservation of amygdalar cell types between humans and rodents (**Figure 2C**). Across all shared homologous genes, we observed substantial correlation between several mouse and human amygdala cell types. For example, our human glutamatergic subcluster Excit.1 (*SLC17A6*+, *SLC17A7*+) most closely correlated with the mouse MeA glutamatergic subcluster ‘N.11’ (Pearson correlation: *r* = 0.455). Indeed the marker genes that were most highly conserved between these subclusters included *SLC30A3, NPTX1*, and *NRN1*. Another notable pair of clusters conserved between species was human inhibitory neuronal subcluster, Inhib.5, and mouse inhibitory subcluster MeA ‘N.8’ (*r* = 0.465). The top shared genes between these clusters included *NPFFR2, GRM8*, and *FOXP2*. Though we observed selective co-expression of *NPFFR2* and *TLL1* in human Inhib.5, we note absence of orthologous *Tll1* expression in all mouse MeA neuronal subclusters (**Figure S6**), including the corresponding cluster ‘N.8’, suggesting species differences in the molecular characteristics of neuronal subpopulations. Importantly, several neuronal subpopulations in the mouse and human datasets lacked strong correlation with each other (e.g. human Excit.2, mouse ‘N.3’ and N.7’), either suggesting possible divergence between species, or unique differences between the cell-type makeup of amygdalar subregions, such that all subpopulations may not be fully represented in our human amygdala sample compared to mouse MeA samples. Our crossspecies analysis demonstrates the conservation of neuronal subtypes between human amygdala and mouse MeA, but highlights potential differences in the cellular distribution and transcriptomic profiles across neuronal subtypes.

### Convergent cell classes with unique molecular signatures across brain regions

We lastly analyzed our 12 homogenate samples (a total of 34,005 nuclei) together at the pan-brain level to identify broad patterns of transcriptional dynamics across the human brain, assessing intra- and inter-regional similarities. We applied the same clustering, annotation, and marker-defining strategies as in region-specific analyses, allowing us to compare the higher-resolution cell type/subpopulation annotation, within each region, to clustering and annotations performed across brain regions. This analysis yielded 17 robust cell type clusters broadly classified into excitatory (Excit) and inhibitory (Inhib) neurons, oligodendrocytes (Oligo), oligodendrocyte precursor cells (OPC), astrocytes (Astro), and microglia (Micro; **Figure S7**). These pan-brain clusters showed some apparent regional specificity (**Figure 3A; Table S5**). For example, excitatory neuronal subclusters Excit.1, Excit.2, Excit.5, and Excit.7 were nearly entirely comprised of nuclei from both cortical regions (DLPFC and sACC), likely reflecting the similar laminar distribution of neuronal subpopulations in the cerebral cortex. Alternatively, Excit.6 was driven by HPC nuclei, whereas Excit.8 had nuclei from both HPC and AMY. Ultimately, these 17 clusters defined across our five regions, analyzed together, provide only a broad overview of similarities between nuclei in a region-agnostic approach, as opposed to better-resolved regionally-defined populations (see Discussion).

**Figure 3:**
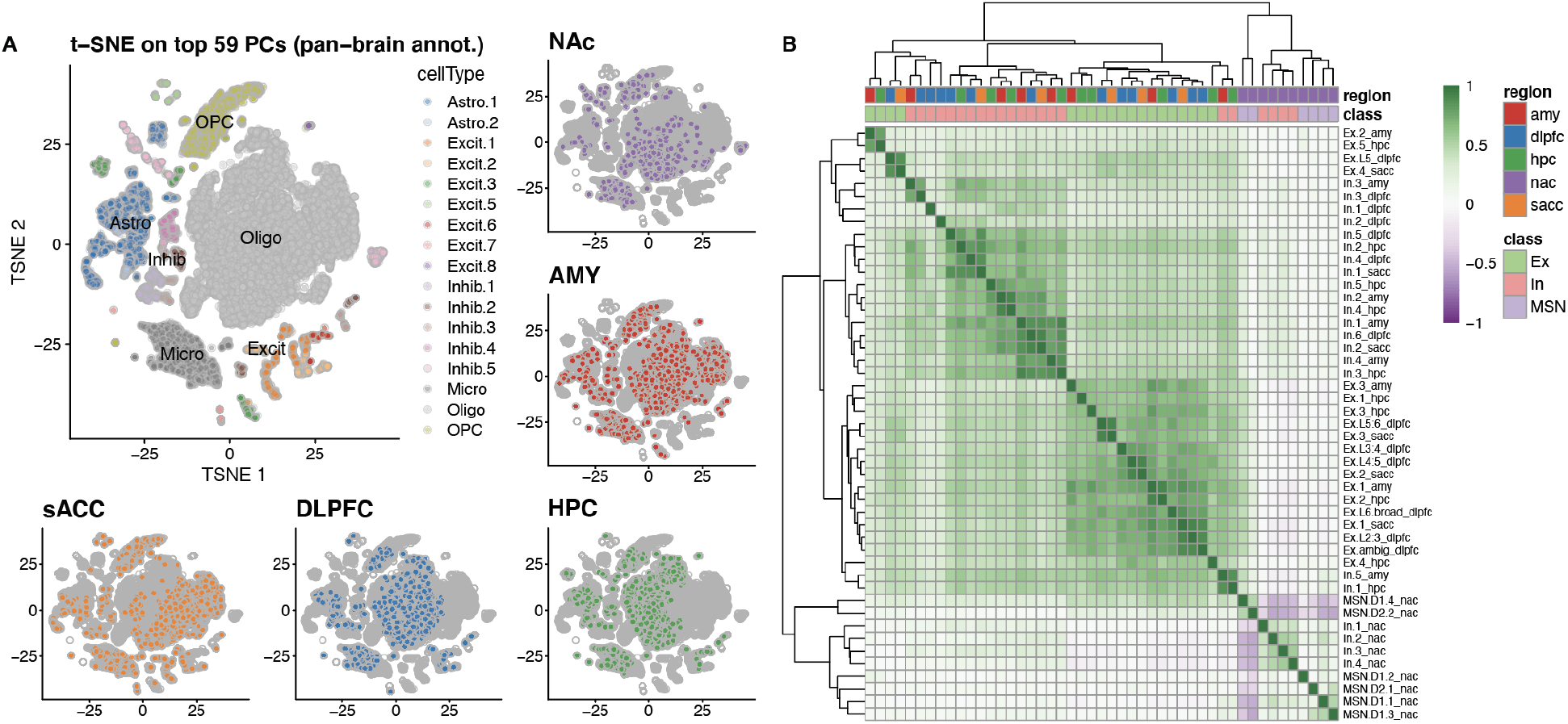
Pan-brain analyses reveal whole brain transcriptomic architecture and neuronal subtype similarities across regions. **(A)** tSNE array of a total of 34,005 nuclei, profiled and clustered across the pan-brain analysis, including their coordinates displayed by each brain region. **(B)** Pairwise correlation of population-defined *t*-statistics, comparing 26,888 genes across a total of 47 neuronal subpopulations, collectively defined across each region (labeled in the suffix). Scale values are of Pearson correlation coefficient.

To complement the subcluster populations described in the previous sections for the NAc and AMY, we additionally defined the catalog of cell type clusters and cluster-specific genes within all other brain regions (sACC, DLPFC, and HPC), separately (**Figure S7**), applying methods using spatial transcriptomics data (Maynard et al., 2020a) to provide layer-specific annotations to our many excitatory neuronal subpopulations for the DLPFC (see Methods). We further benchmarked our transcriptomic profiles against other published datasets that profiled similar regions in the postmortem human brain. Overall, our HPC subpopulations correlated well with the broad cell classes as reported in (Habib et al., 2017); **Figure S8**). We additionally observed strong overlap between our DLPFC to the reported PFC profiles from (Velmeshev et al., 2019); **Figure S9**), or similarly, sACC to the ACC set (**Figure S10**). Interestingly, our sACC subpopulations did not correlate more strongly with the ACC subpopulation profiles than their corresponding PFC profiles from (Velmeshev et al., 2019), and our DLPFC subclusters generally correlated only slightly more strongly to PFC than ACC subpopulations. This suggests that these cortical regions share a high degree of overlap in their nuclear transcriptomic profiles. The strength of correlation to these benchmark datasets demonstrates the robustness and utility of our pipeline, and the presented data significantly expand the existing repository of postmortem brain snRNA-seq datasets.

We next compared all regionally-defined subcluster expression patterns to generate a global view of the transcriptomic architecture across the five brain regions. Each glial cell subpopulation (Oligos, Astros, OPCs, and Micros) had highly consistent gene expression patterns across all five brain regions, suggesting a lack of overall regional specificity (**Figure S11**), in line with previous analyses of broad non-neuronal cell populations using DNA methylation data (Rizzardi et al., 2019). Within the neuronal set of region-specific annotations, totaling 47 neuronal subpopulations, most inhibitory or excitatory populations preferentially expressed genes that clustered these classes together across brain regions (**Figure 3B**). Notably, all neuronal subtypes in the NAc (interneuron or MSNs) correlated poorly or less-strongly with either excitatory or inhibitory subtypes from all other regions, reflecting divergent nuclear RNA expression profiles and the unique functional specializations of the neuronal populations in this brain region. These comparisons also suggested a potential excitatory function in the NAc-specific MSN.D1.4 subcluster, as this profile correlated more strongly with excitatory subclusters across other brain regions than inhibitory subclusters. This particular D1 MSN subcluster most highly correlated with one D2 MSN subcluster (MSN.D2.2; Pearson correlation: *r* = 0.50), reflecting their sharing of top marker genes (**Figure S4**), and strikingly these two NAc subpopulations negatively correlate with all other MSN or inhibitory interneuron populations within the NAc, further highlighting the specializations across NAc neuronal subpopulations. We also observed strong similarities between unique pairs of neuronal subpopulations across regions, such as between AMY and DLPFC (‘In.3_amy’ and ‘In.3_dlpfc’; *r* = 0.74). Indeed, this AMY inhibitory subpopulation shares many top markers with its DLPFC counterpart (**Table S4**), such as *GABRD* and *MME*, the latter encoding for the neuropeptide-cleaving protein, Neprilysin. Further, we observed analogous pairings between each of layer-specific excitatory DLPFC and spatially-undefined sACC subpopulations (e.g. ‘Ex.L5_dlpfc’ and ‘Ex.4_sacc’; *r* = 0.86). This suggests that spatially-registered snRNA-seq information, as generated in the DLPFC (Maynard et al., 2020a), can be projected into regions where cell type architecture is expected to be similar. Such an approach is useful since spatial transcriptomic data generation has not yet reached the pace of sc/sn-RNAseq technologies, and remains unavailable for most brain regions.

### Enrichment of region-specific cell subtypes in psychiatric disease and substance use

Genome-wide association studies (GWAS) have identified a plethora of genetic risk variants or loci (segregating variants in linkage disequilibrium, or LD) for common psychiatric disorders, including schizophrenia (SCZ: (Pardiñas et al., 2018; Schizophrenia Working Group of the Psychiatric Genomics Consortium, 2014)), autism spectrum disorder (ASD: (Grove et al., 2019)), bipolar disorder (BIP: (Stahl et al., 2019)), major depressive disorder (MDD: (Howard et al., 2019; Wray et al., 2018)), and posttraumatic stress disorder (PTSD: (Nievergelt et al., 2019)). Additionally, a large GWAS was recently performed with 1.2 million individuals to identify the genetic risk and correlations for alcohol and tobacco use (Liu et al., 2019). Approaches have been developed to identify the biological context or relevance of the hundreds of risk loci that are often identified for a given disorder or phenotype, such as LD score regression (Finucane et al., 2015), which assesses the heritability of complex phenotypes across input categories/genomic regions and their measured LD with single nucleotide polymorphism (SNP)-level variants. Multi-marker Analysis of GenoMic Annotation (MAGMA) (de Leeuw et al., 2015) is an alternative approach that defines gene-level localization of GWAS risk, then integrates this with gene set observations, affording flexibility to assess a variety of marker lists, such as for our brain region-specific snRNA-seq subcluster profiles, in two separable analyses.

We used MAGMA to identify which cell subtypes in this study harbored aggregated genetic risk for psychiatric disorders, and found robust signals across many nuclear profiles in each of the five profiled brain regions. As expected, many DLPFC and HPC neuronal subtypes exhibited significant effect sizes for both SCZ and BIP GWAS, spanning both excitatory and inhibitory subpopulations (**Figure S12**), extending and strengthening previous findings in (Bryois et al., 2020; Skene et al., 2018). However significant association signals with BIP genetic risk only remain in the excitatory, layer-specific DLPFC subpopulations, after controlling with the strict Bonferroni multiple test correction across all MAGMA gene set tests (threshold *p*-value < 6.8e-5). Conversely, subpopulations of both excitatory and inhibitory classes associated with BIP risk at this threshold in HPC. This was additionally the case for sACC subclusters, which suggests potential regional differences in inhibitory subpopulations between the two cortical brain regions, in their manifestation of genetic risk for bipolar disorder. None of the subcluster profiles in these cortical or hippocampal regions retained significant signal for aggregated risk for GWAS in ASD, MDD, or PTSD, after Bonferroni correction, though there were some region subpopulations with less-stringent FDR-significant signals (controlling for false discovery rate < 0.05) across all tests for these disorders (**Table S6**).

Previously, it has been shown that broad mouse striatal neuronal populations (interneurons, Drd1, and Drd2-expressing medium spiny neurons, or MSNs) additionally associated with SCZ (Skene et al., 2018) and BIP (Bryois et al., 2020) genetic risk. We demonstrate that some of our refined subpopulations in the human NAc, including MSN.D1.4, and MSN.D2.1, and D2.2 exhibit strong associations to schizophrenia with variable effect sizes, after Bonferroni correction (**Figure 4A**). MSN.D1.3 and MSN.D2.1 both additionally associated with BIP at this threshold, and interestingly, D1.3 was the strongest-associating subpopulation to BIP, whereas D2.1 had the strongest association to SCZ. Within the amygdala, we observed significant associations in each of our neuronal subpopulations to SCZ (**Figure 4B**). Notably, the strongest signal and effect sizes were observed across each of Inhib.1 through Inhib.5, in comparison to AMY excitatory subpopulations, and Inhib.4 exhibited the strongest effect size across all regionally-defined subpopulations tested in any GWAS phenotype (*β* = 0.21). Thus, our analysis here with the NAc show not only complementary findings of Drd1 and Drd2-expressing striatal MSN associations with schizophrenia and bipolar disorder, but we dissect these mouse association signals with more relevant human interneuron and MSN subpopulations, and further extend this analysis to human AMY snRNA-seq-defined subpopulations.

**Figure 4.**
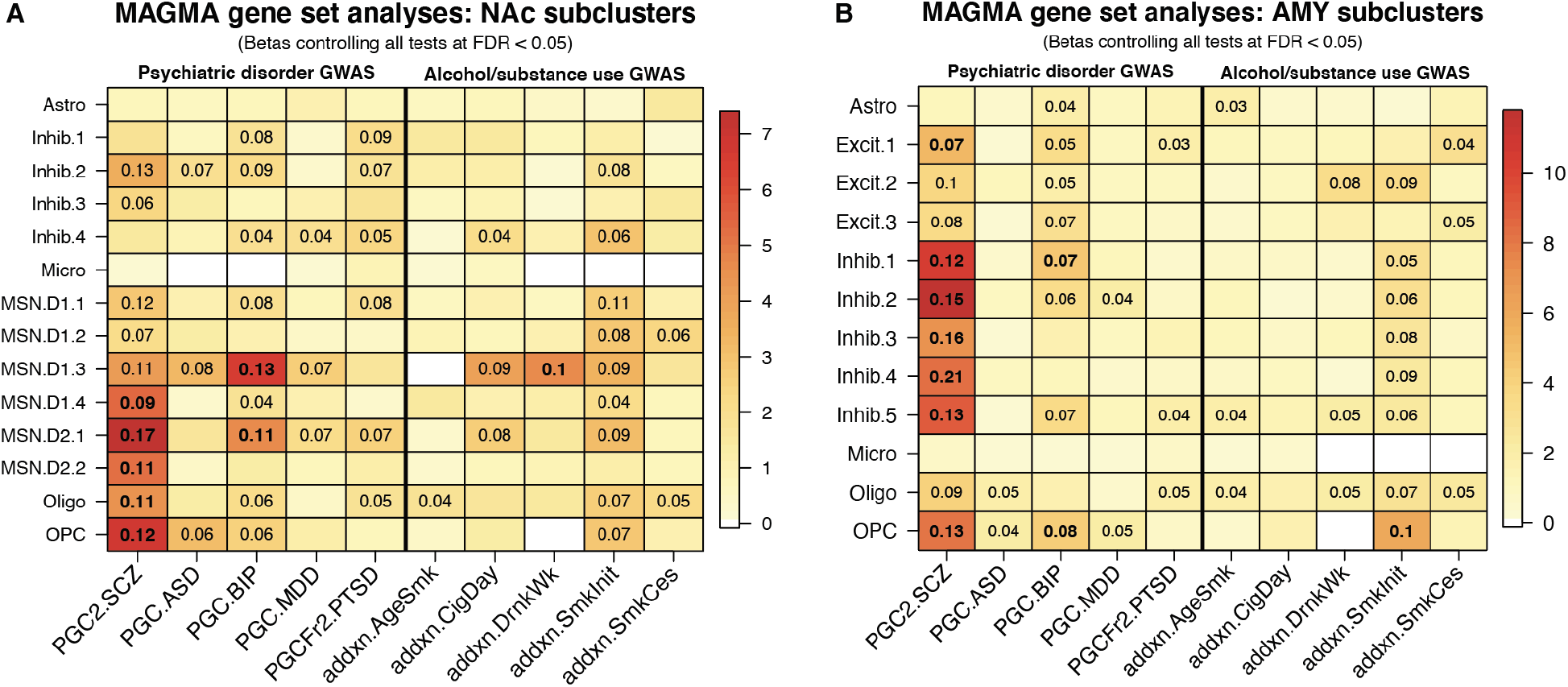
Genetic associations of NAc and AMY cell populations with psychiatric disease and addiction phenotypes. **(A)** MAGMA associations for each of 14 subpopulations profiled in human NAc or **(B)** 12 subpopulations profiled in human AMY. Heatmap is colored by empirical −log10(*p-value*) for each association test. Displayed numbers are the effect size (***β***) for significant associations (controlled for false discovery rate, FDR < 0.05), on a *Z* (standard normal) distribution. Bolded numbers are those that additionally satisfy a strict Bonferroni correction threshold of *p* < 6.8e-5.

We further tested for alcohol and tobacco use GWAS (Liu et al., 2019) genetic risk associations across subcluster profiles from each of our brain regions, focusing on the subcortical regions centered in reward circuitry, the NAc and AMY, and their subcluster profiles described above. This highlighted various MSN and inhibitory subpopulations in the NAc associating with genetic risk for regular smoking behavior (‘SmkInit’) at FDR < 0.05 (**Figure 4A**), along with associations to multiple AMY excitatory and inhibitory neuronal subpopulations (**Figure 4B**). Additionally, both Oligo and OPC clusters from both regions exhibited aggregated risk for regular smoking behavior. However after applying the above Bonferroni threshold across all regions and phenotypes tested, only the AMY ‘OPC’ population retained significant association with this behavior, whereas no NAc subpopulations met this significance threshold. Conversely, only the NAc contained a subpopulation harboring Bonferroni-significant risk for any of the other phenotypes in the addiction GWAS. This was the *GABRQ* and *TAC1*-expressing subcluster MSN.D1.3, which associated with heaviness of drinking (‘DrinkWk’). Collectively, these results provide complementary human findings for genetic risk associations to those previously described for psychiatric disease, further identifying subpopulations in the NAc and AMY harboring aggregated risk for substance use behaviors.

## Discussion

In this study, we used snRNA-seq to profile five human brain regions across the ventral striatum (NAc), limbic system (AMY and HPC), and two cortical subregions (sACC, DLPFC). While single-nucleus transcriptomic profiling in the postmortem human brain has rapidly accelerated, most efforts to date (Mathys et al., 2019; Nagy et al., 2020; Velmeshev et al., 2019) have focused on cortical regions and the hippocampus. This is the first study, to our knowledge, to systematically profile and compare across additional human brain areas selected for their function associated with risk for neuropsychiatric disorders and addiction. Specifically, we focused on the NAc and the AMY given their roles in emotional processing and reward signaling. While this study was performed only among neurotypical donors, the strong cell typespecific associations to genetic risk for these disorders provide important information about disease etiology. This link to genetic risk is important, given that differential gene expression identified in case-control studies of postmortem tissue are difficult to interpret and may represent consequences (and not causes) of these disorders (Collado-Torres et al., 2019; Jaffe et al., 2020). More generally, understanding the transcriptomic architecture and cell type composition across the normal human brain is crucial to understanding the etiology of disease, the molecular pathology observed in postmortem tissues, and novel potential disease targets. Our study is a significant contribution as it demonstrates differential enrichment of disease risk in snRNA-seq-defined cell populations across multiple brain regions, including the NAc and AMY, which have not yet been profiled at the single-nucleus level in the human brain.

The NAc is a central hub for reward signaling, and altered function in circuits encompassing the NAc is implicated in a number of psychiatric disorders as well as drug addiction. Hence, we sought to define molecular profiles for NAc cell types, with a specific focus on functionally dichotomous subtypes of DRD1- and DRD2-expressing MSNs. Consistent with studies that used single-cell sequencing to profile the mouse striatum, including ~1000 striatal cells in each study (Gokce et al., 2016; Stanley et al., 2020), we identified several discrete subpopulations of D1 and D2-expressing MSNs in human NAc. Similar to Gokce et al., we also identified 2 discrete subpopulations of D2 MSNS, including a subtype expressing the serotonin receptor gene *HTR7*. However, distinct from Gokce et al. and Stanley et al., we identified 4 discrete subpopulations of D1-MSNs, which we validated by smFISH. Several reasons may explain why we identified different discrete D1 cell types, including differences in species (human vs. mouse), region (NAc-specific vs. striatum-wide), sample preparation (whole cells vs. nuclei), number of MSNs profiled (about 8x greater in our dataset) and single cell technology employed (10x Genomics Chromium vs. SMART-Seq v2). However, in agreement with these studies, we also observed co-expression of *DRD1* and *DRD2* in a small subset of MSNs. While these dual-expressing neurons did not emerge as their own cluster, they were largely found in the D1.3 subpopulation (**Figure 1B**). Interestingly, this cluster showed the strongest enrichment of genes associated with psychiatric disease and addiction, indicating that this particular subpopulation might be especially vulnerable to dysfunction in these disorders. Indeed, among D1 subtypes, D1.3 MSNs show selective expression of *CRHR2*, a gene encoding corticotropin releasing hormone receptor 2, suggesting that they may be particularly susceptible to the effects of corticotropin-releasing hormone (CRH), which is released and mediates the physiological and behavioral response to stress, modulating several neurotransmitter systems, including dopamine release (Bonfiglio et al., 2011; Payer et al., 2017). Given that dysfunction of the CRH system has been associated with many psychiatric disorders, including depression, anxiety, and PTSD (Claes, 2004), understanding which cell types express CRH receptors may aid in more specific targeting of the stress axis for therapeutic developments.

Similar to Gokce et al., we also observed promiscuous expression of “typical” D1 and D2 neuropeptide marker genes (*TAC1* and *PENK*, respectively) in both D1 and D2 MSN subpopulations, providing further evidence that these classic markers may not be as selectively expressed as previously understood. Future studies using spatial transcriptomic approaches will be important to clarify whether *TAC1*-expressing D1 and D2 MSN subpopulations show topographical organization in the NAc core vs. shell. Anatomical location may explain differences in *TAC1* and *PENK* expression in specific MSN subpopulations, as it is well established that specific neuropeptides are expressed in a spatial gradient across the core and shell (Prensa et al., 2003; Salgado and Kaplitt, 2015; Stanley et al., 2020; Voorn et al., 1989). To better interpret clinical implications of studies that focus on circuitry encompassing the NAc in animal models, further understanding of similarities and differences across species for cell types that contribute to NAc function are important. While many cell populations were conserved between rat and human NAc (Savell et al., 2020), we did observe differences in specific MSN subpopulations, which may indicate unique molecular features between analogous MSN subpopulations and/or the existence of divergent MSN subclasses, as exemplified by the lack of a corresponding human MSN subpopulations with the rat ‘*Grm8-MSN*’ subpopulation (**Figure 1E**). However, given the small positive correlations measured with human D1.1 (*r* = 0.19) and D1.3 (*r* = 0.26) subtypes, it is possible that this *Grm8*-expressing population encompasses the species-equivalent of these less abundant D1 subtypes, since D1.1 expresses abundant *GRM8*, while D1.3 expresses little (**Figure S4**), even though the latter correlates more strongly with the rat ‘*Grm8-MSN*’. We also were unable to identify a population of cholinergic interneurons. While cholinergic interneurons are thought to be more abundant in the human neostriatum compared to the rodent, where they only account for ~0.3% of neurons (Graveland et al., 1985; Rymar et al., 2004; Tepper and Bolam, 2004), we think it is likely that the low rate of sampling and this population’s relative rarity accounts for this lack of observation.

In addition to profiling NAc cell types, we also generated a molecular taxonomy of human amygdala cell types. We identified eight distinct neuronal subpopulations as well as accompanying gene marker annotations, including *NRN1* (neuritin) and *NPTX1* (neuronal pentraxin 1) for the AMY Excit.1 subcluster. Neuritin is a neurotrophic factor which modulates neurite outgrowth and plasticity (Yao et al., 2018), whereas neuronal pentraxin 1 regulates neuron excitability via synapse density (Figueiro-Silva et al., 2015). Additionally, the highest *SLC17A6* (VGLUT2)-expressing subcluster, Excit.2, specifically expressed high levels of *VCAN* (Versican) amongst other neuronal subpopulations, which has multiple isoforms exhibiting different mechanisms for synaptic regulation (Horii-Hayashi et al., 2008). Among the diverse set of inhibitory subpopulations in the AMY, the stress modulator *CRH* was specifically enriched in Inhib.3 and Inhib.4. Top markers in the AMY Inhib.5 subcluster included *NPFFR2* (Neuropeptide FF Receptor 2) and *TLL1* (Tolloid-Like 1), which are both associated with glucocorticoid signaling and the response to stress (Lin et al., 2016, 2017; Tamura et al., 2005). Interestingly, *TLL1* harbors top intronic genetic variant associations for PTSD in multiple European American cohorts (Xie et al., 2013), suggesting that dysfunction of neurons in Inhib.5 subcluster might mediate genetic risk for PTSD. Using MAGMA, we assessed whether the top nuclear marker genes for AMY subclusters associated with PTSD genetic risk (Nievergelt et al., 2019), and found that Inhib.5 was indeed one of two AMY neuronal subclusters that reached significance (FDR < 0.05) across all gene set tests (**Figure 4B**). This result did not maintain significance using a stricter Bonferroni threshold; however this GWAS is underpowered, and it is anticipated that with increased sample size in future iterations, stronger association signals for PTSD genetic risk may be revealed. Comparing human AMY subcluster profiles to data from the mouse medial amygdala (MeA; (Chen et al., 2019)), we found that Inhib.5 and its corresponding population in mouse (MeA ‘N.8’ subcluster, **Figure 2C**) were the most strongly correlated neuronal subpopulations. While *Tll1* expression was notably absent in mouse MeA, *Npffr2* and other top MeA ‘N.8’ marker genes were shared with Inhib.5 (**Figure S6**). These insights highlight the importance of deriving reference snRNA-seq datasets across the human brain, as molecular gene markers may not be shared across species between analogous neuronal subpopulations.

We next used the snRNA-seq data from the five profiled regions to ask whether identified cell subpopulations harbored aggregate genetic risk for various neuropsychiatric disorders and/or features of substance use. We confirmed previous findings by identifying strong associations for neuronal subpopulations in the DLPFC and HPC with both schizophrenia (SCZ) and bipolar disorder (BIP) (Bryois et al., 2020; Skene et al., 2018), and significantly extended these findings by providing associations with specific sACC excitatory and inhibitory populations (**Figure S12**). Additionally, we not only confirmed previously observed associations to broad striatal populations defined in the mouse, but used our human NAc subcluster profiles to further refine these findings by demonstrating that SCZ most strongly associated with subpopulation MSN.D2.1, whereas D1.3 had the strongest association for BIP (**Figure 4A**). This suggests, for the first time, that individual populations of dopaminoceptive (DRD1/2) neurons in the human NAc may be differentially associated with SCZ and BIP. We also found that specific subpopulations of inhibitory interneurons in the human AMY were preferentially associated with SCZ, with AMY Inhib.4 exhibiting the strongest effect across the five profiled brain regions. These observations highlight a potential role for these subcortical brain regions in mediating genetic risk for SCZ and BIP.

As both the NAc and AMY play critical roles in reward signaling, we also evaluated enrichment of genetic risk for addiction or substance use behaviors (Liu et al., 2019). Intriguingly, the genetic risk for adopting regular smoking associated more broadly across most neuronal populations, whereas other phenotypes assessed in this addiction GWAS showed more preferential associations to certain subpopulations. This suggests that the risk for adopting addictive-like behaviors might affect these brain regions more broadly than specific features of addiction (**Figure 4A/B**). With regard to the other features, the MSN.D1.3 subpopulation significantly associated with genetic risk for heaviness of drinking (‘DrnkWk’) and smoking (‘CigDay’), after Bonferroni and FDR correction, respectively. As a top marker for this subpopulation was *CRHR2*, this might be a key population in understanding these features of addiction. Indeed, many rodent studies have implicated CRH receptors in alcohol consumption and alcohol dependence (Heilig and Koob, 2007; Yong et al., 2014). Finally, though no neuronal AMY subpopulations met our strict Bonferroni threshold for association, the potentially HPA axis-involved ‘Inhib.5’ population exhibited more FDR-significant associations to addiction phenotypes in this GWAS than any other neuronal subpopulation, suggesting that this *NPFFR2/TLL1*-expressing AMY subpopulation might be of interest in understanding the reward circuitry underlying substance use. From these analyses, we surveyed our diversity of neuronal subpopulations profiled in the NAc and AMY for their clinical relevance in psychiatric disease and addiction behaviors. Additionally, we have extended such analyses for these regions, which have formerly only been performed on cell-type profiles defined in murine models (Bryois et al., 2020; Skene et al., 2018) to their relevant human context, and with increased resolution of molecularly-defined subpopulations. Finally, we narrowed down on those subpopulations manifesting the greatest genetic risk, potentially highlighting some neuronal subclasses mediating certain substance use behaviors.

Future studies conducted in multiple brain regions warrant important considerations in study and analysis design for interpretability. Here, we performed: (1) a ‘pan-brain’ analysis with all of our 12 homogenate (non-NeuN-selected) samples, combined in a region-agnostic manner (**Figure 3A)**; and contrasted this against (2) performing the above cell type clustering and analysis within each of the regions, separately. There were few benefits of the pan-brain approach, as identified clusters showed little regional variation (e.g. glial and broad neuronal populations) or near-complete segregation by brain region (**Table S5**). We therefore recommend prioritizing region-specific analyses even when data from multiple brain regions are collected, and then integrating information from these analyses across brain regions, over performing combined-region analyses. Of course, the optimal approach may depend on the research question being asked.

While we identified and characterized a diversity of robust neuronal subpopulations with our analytical pipelines for this study, we recognize that our sample sizes will not capture all cell types or subpopulations--even for our most-sampled brain region, the NAc, such as cholinergic interneurons mentioned above. The most direct evidence for this is that there remains some bias in donor makeup of certain subpopulations (**Table S3**). However, despite steps to mitigate the impact of the small input for our sample processing protocol (see Methods), we expect some degree of sampling bias since cell type makeup is not expected to be homogeneous within a single region. For example, the NAc core or shell have different functional properties, and differ in regards to their afferent and efferent connections, and thus differences in cell composition across these two subregions is expected (Heimer et al., 1991; Li et al., 2018b; Zahm and Heimer, 1993). Integration of spatial transcriptomic technologies with snRNA-seq data in these regions (Maynard et al., 2020a) will help resolve expected heterogeneity across these adjacent subregions. To similar effect, we acknowledge that we may have missed some expected nonneuronal, or potentially even neuronal, subpopulations in our approaches to define these subcluster profiles (see Methods). For example, we expected some number of endothelial cells and pericytes in each of our regions, making up the vasculature of the brain; however, these likely clustered with the more abundant microglia, due to a shared cell lineage (Bailey et al., 2006). Ultimately, our goal was to better characterize neuronal subpopulations across these five brain regions, and with these sample sizes being still limited, we did not pursue any nested or further cluster refinement of the presented populations. With all of this in mind, we were able to validate the presence and relative ratios of subpopulations defined by our snRNA-seq analytical pipeline via smFISH.

Another caveat to these snRNA-seq data is the lack of gene expression information from the cytosolic compartment, such as the neuropil. This is an important caveat given that synaptic signaling is implicated in neuropsychiatric disorders, and gene products localized to the synapse are enriched for SCZ genetic risk (Skene et al., 2018). In addition, mRNA from some expected marker genes, e.g. *PVALB*, may be preferentially localized to the cytosol, as demonstrated with smFISH for the *GAD1*+ interneuron ‘Inhib.3’ population in the NAc (**Figure S5**). However, this seems to be cell population-specific, as *PVALB* was highly expressed in some DLPFC subpopulations (data not shown; see *Data and Software Availability*). These and observations by others thus emphasize that snRNA-seq will not capture the full transcriptomic profile of cell populations, including activation-induced or disease-associated molecular changes restricted to the cytosol (Thrupp et al., 2020). However, as we have previously demonstrated (Maynard et al., 2020a), snRNA-seq-defined cell populations can be registered to spatial transcriptomic data, which does retain such information, for further characterization of transcriptomic profiles.

In summary, we used snRNA-seq to profile five human brain regions. We defined transcriptomic profiles for 68 regionally-defined cell type subpopulations and characterized the architecture of molecular relationships across brain regions. We finally identified associations with genetic risk for neuropsychiatric disorders and addiction in unique neuronal subpopulations in the NAc and AMY. This study takes a large step towards mapping the single-nucleus transcriptomic atlas of the human brain, further demonstrating such a utility in understanding the diversity of cell populations and their roles in biology and disease.

## Methods

### Post-mortem human tissue

Post-mortem human brain tissue from five neurotypical donors of European ancestry from age 40 to 62 (**Table S1**) was obtained by autopsy from the Office of the Chief Medical Examiner for the State of Maryland under State of Maryland Department of Health and Mental Hygiene Protocol 12-24. Clinical characterization, diagnoses, and macro- and micro-scopic neuropathological examinations were performed on all samples using a standardized paradigm, and subjects with evidence of macro- or micro-scopic neuropathology were excluded. Details of tissue acquisition, handling, processing, dissection, clinical characterization, diagnoses, neuropathological examinations, RNA extraction and quality control measures have been described previously (Lipska et al., 2006). Dorsolateral prefrontal cortex (DLPFC) and hippocampus (HPC) tissue was microdissected using a hand-held dental drill as previously described (Collado-Torres et al., 2019). The subgenual Anterior Cingulate Cortex (sACC) was dissected under visual guidance from the medial aspect of the forebrain at the level of the rostrum of the corpus callosum. Dissections were performed ventral to the corpus callosum, and dorsal to the orbital frontal cortex (BA11). Medially it was bounded by the interhemispheric fissure, while laterally it was bounded by the corona radiata/centrum semiovale. For the amygdala, a block containing the structure was dissected under visual guidance at the level of its maximal size, taken from a 1 cm thick slab of one hemisphere, and sectioned in the coronal plane. The amygdala block was chosen by visual inspection at a level that contained the maximal number of subnuclei. Landmarks for selection of the amygdala block included presence of the internal and external segments of the globus pallidus, the anterior commissure, and optic tract. The block containing the nucleus accumbens was taken from a 1 cm thick slab of one hemisphere, and sectioned in the coronal plane. The NAc block was chosen at a level where the putamen and caudate are joined by the accumbens at the ventral aspect of the striatum, with clear striations separating the putamen from the caudate. Additional landmarks include the presence of the anterior aspect of the temporal lobe and the claustrum.

### snRNAseq data generation

We performed single-nucleus RNA-seq (snRNA-seq) on 14 samples from five individual donors (*n*=2 DLPFC, *n*=3 HPC, *n*=2 AMY, *n*=2 sACC, *n*=5 NAc) using 10x Genomics Chromium Single Cell Gene Expression V3 technology (Zheng et al., 2017). Nuclei were isolated using a “Frankenstein” nuclei isolation protocol developed by Martelotto *et al*. for frozen tissues (Habib et al., 2016, 2017; Hu et al., 2017; Lacar et al., 2016; Lake et al., 2016). Briefly, ~40mg of frozen, ground tissue was homogenized in chilled Nuclei EZ Lysis Buffer (MilliporeSigma #NUC101) using a glass dounce with ~15 strokes per pestle. Homogenate was filtered using a 70μm-strainer mesh and centrifuged at 500 x g for 5 minutes at 4°C in a benchtop centrifuge. Nuclei were resuspended in the EZ lysis buffer, centrifuged again, and equilibrated to nuclei wash/resuspension buffer (1x PBS, 1% BSA, 0.2U/μL RNase Inhibitor). Nuclei were washed and centrifuged in this nuclei wash/resuspension buffer three times, before labeling with DAPI (10μg/mL). For 2 NAc samples from individual donors, nuclei were additionally labeled with Alex Fluor 488-conjugated anti-NeuN (MilliporeSigma cat. #MAB377X), at 1:1000 in the same wash/resuspension buffer to facilitate enrichment of neurons during fluorescent activated cell sorting (FACS). Samples were then filtered through a 35μm-cell strainer and sorted on a BD FACS Aria II Flow Cytometer (Becton Dickinson) at the Johns Hopkins University Sidney Kimmel Comprehensive Cancer Center (SKCCC) Flow Cytometry Core into 10X Genomics reverse transcription reagents. Gating criteria hierarchically selected for whole, singlet nuclei (by forward/side scatter), G0/G1 nuclei (by DAPI fluorescence), and NeuN-positive cells for the respective NeuN-enriched samples. A “null” sort of nuclei into the wash buffer was additionally performed from the same preparation, for quantification of nuclei concentration and to ensure that sorted nuclei were intact and free of debris. For each sample, approximately 8,500 single nuclei were sorted directly into 25.1μL of reverse transcription reagents from the 10x Genomics Single Cell 3’ Reagents kit (without enzyme). Libraries were prepared according to manufacturer’s instructions (10x Genomics) and sequenced on the Next-seq (Illumina) at the Johns Hopkins University Transcriptomics and Deep Sequencing Core.

### snRNAseq raw data processing

We processed the sequencing data with the 10x Genomics’ Cell Ranger pipeline, aligning to the human reference genome GRCh38, with a reconfigured GTF such that intronic alignments were additionally counted given the nuclear context, to generate UMI/feature-barcode matrices (https://support.10xgenomics.com/single-cell-gene-expression/software/pipelines/latest/advanced/references). Per the output metrics of Cell Ranger, each sample was sequenced to a median depth of 253.0M reads (IQR: 148.7M-274.9M). We started with raw feature-barcode matrices from this output for analysis with the Bioconductor suite of R packages for single-cell RNA-seq analysis (Amezquita et al., 2020) using Bioconductor (Huber et al., 2015) versions 3.10 and 3.11. For quality control (QC) and nuclei calling, we first used a Monte Carlo simulation-based approach to assess and exclude empty droplets or those with random ambient transcriptional noise, such as from debris (Griffiths et al., 2018; Lun et al., 2019). This was then followed by mitochondrial rate adaptive thresholding, which, though expected to be near-zero in this nuclear context, we applied a 3x median absolute deviation (MAD) threshold, to allow for flexibility in output/purity of nuclear enrichment by FACS using *scater*’s `isOutlier` (Lun et al., 2016). This QC pipeline yielded 5,399 high-quality nuclei from the DLPFC, 10,444 nuclei from HPC, 6,632 nuclei from AMY, 7,047 nuclei from sACC, and 13,241 nuclei from NAc. Collectively, these exhibited a median unique molecular identifier (UMI) count of 8,747 (interquartile range, IQR: 5,280-19,895 UMIs) per nucleus, and a median detected gene count of 3,047 (IQR: 2,224-5,359) genes captured per nucleus. These feature-barcode gene counts were then rescaled across all nuclear libraries, using *scater*’s `librarySizeFactors` (Lun et al., 2016). Finally, these rescaled counts were log_2_-transformed for identification of highly-variable genes (HVGs) with *scran*’s `modelGeneVar` (Lun et al., 2016), taking all genes with a greater variance than the fitted trend.

### Dimensionality reduction and clustering

Principal components analysis (PCA) was then performed on the HVGs to reduce the high dimensionality of nuclear transcriptomic data, both in region-specific analyses and pan-brain analyses. The optimal principal component (PC) space was defined with iterative graphbased clustering to determine the *d* PCs where resulting *n* clusters stabilize, with the constraint that *n* clusters </= (*d* + 1) PCs (Lun et al., 2016), resulting in a chosen *d* between 45-96 PCs, in the region-specific analyses, or 204 PCs for the pan-brain analysis (all homogenate, DAPI-sorted samples). In this PCA-reduced space, graph-based clustering was performed to identify what we classified as preliminary clusters; specifically, *k*-nearest neighbors with *k*=20 neighbors and the Walktrap method from R package *igraph* (Csardi and Nepusz, 2006) for community detection. We used an increased *k* neighbors (from the default *k*=10) as a means to increase the connectivity in the *k*NN graph, as an alternate approach for handling potentially donor-driven preliminary clusters, instead of manually identifying batch-correlated PCs and removing these. We then took all feature counts for these assignments and pseudo-bulked counts (Crowell et al., 2019; Kang et al., 2018; Lun and Marioni, 2017) across these preliminary nuclear clusters, rescaling for combined library size and log-transformed normalized counts, as above. With the pseudo-bulked count profiles, we then performed hierarchical clustering to identify preliminary cluster relationships, and finally merged with the `cutreeDynamic` function of R package *dynamicTreeCut* (Langfelder et al., 2016), or keeping split clusters at the preliminary resolution, if generally well-represented across donors, as this suggested biologically valid subpopulations (for example, neuronal subtypes) as opposed to more likely batch-driven preliminary clusters. However, in some cases, cluster marker identification (see below) suggested sample bias in true, biological subpopulations (see Discussion). The final clusters merged at the appropriate tree height were then annotated for broad cell type identity with well-established cell type markers (Mathys et al., 2019), and with a numeric suffix where multiple broad cell class populations were defined (‘*Excit.1*’, ‘*Excit.2*’, etc.). We also used Bioconductor package *scater*’s (McCarthy et al., 2017) implementation of non-linear dimensionality reduction techniques, *t*-SNE (van der Maaten and Hinton, 2008) and UMAP (McInnes et al., 2018), with default parameters and within the aforementioned optimal PC space (or a reduced dimensional space in the case of the NAc and AMY, for t-SNE coordinates which would better reflect the final subcluster distribution), simply for visualization of the high-dimensional structure in the data, which complemented the clustering results. Additionally, in each of our five within-region analyses, a small cluster driven by low transcript capture would remain even after hierarchical cluster-merging of preliminary clusters, but these were removed prior to downstream analyses and from the *t*-SNE display, resulting in a final *n* nuclei analyzed per region of: 5,231 from the DLPFC, 10,343 nuclei from HPC, 6,582 nuclei from AMY, 7,004 nuclei from sACC, and 13,148 nuclei from NAc (an average of 98.8% nuclei kept post-QC, above). These final numbers of nuclei analyzed per regionally-defined subcluster and by donor can be found in **Table S3**.

### Cluster marker identification

For marker identification with our final clusters defined in each brain region or at the pan-brain-level analysis, we utilized *scran*’s `findMarkers` (Lun et al., 2016) function for two sets of statistics:

1. Pairwise *t*-tests, to identify differences between each cluster, or
2. Implementing the function `findMarkers` to perform a cluster-vs-all-other-nuclei *t*-test iteration

In both cases, we included a donor/processing date covariate to model (in the `design=`parameter) on these expected and unwanted batch effects. The latter approach, 2), we consider a less-stringent marker test for enriched genes in a given cluster, but which would not necessarily differentiate between said cluster and all others. We used the results from both tests to interpret cell type identity beyond the broad classes (excitatory vs. inhibitory neuron), and to identify markers to probe via smFISH (below). The top 40 markers from each test result are provided for each regionally-defined subpopulation in **Table S4** (regions separated by worksheet), where the ‘*_pw*’ suffix corresponds to the pairwise tests (set *1*), and ‘*_1vAll*’ to the enriched expression test (set *2*).

Importantly, 2) can be used to return a statistic, Cohen’s D, or the standardized log-fold change, which we used to back-compute a single *t*-statistic for each cluster per gene, using:

`t = std.logFC * sqrt(n)`, where `n =` the total *n* nuclei (per region/dataset)
* Back-computing a single *t*-statistic cannot be generated with the result of 1) due to pairwise testing.

These *t*-statistics can finally be used to compare such ‘transcriptomic profiles’ to those we computed for publicly-available datasets, using the provided cell type annotations (or across our 5 regions), and compute the Pearson correlation coefficient, as was done in the spatial registration approaches in *spatialLIBD* (Maynard et al., 2020a). To perform cross-species conservation analyses, we generated these *t*-statistics per gene per reported cell annotation, subsetting on shared homologous genes between our human data and rat or mouse, using the ‘HomoloGene.ID’ identifier provided by (http://www.informatics.jax.org/downloads/reports/HOM_AllOrganism.rpt), before computing the pairwise correlations. In the case of “many-to-many” scenarios, we took the highest-expressing paralog as the surrogate for each homologous pair, though these were small sets of genes in both rat and mouse cases.

### GWAS association analyses with MAGMA

The latest version (*v1.08*) of Multi-marker Analysis of GenoMic Annotation (MAGMA; (de Leeuw et al., 2015) was used to test for genetic risk association of our 68 regionally-defined subpopulations with schizophrenia (SCZ: (Pardiñas et al., 2018; Schizophrenia Working Group of the Psychiatric Genomics Consortium, 2014)), autism spectrum disorder (ASD: (Grove et al., 2019)), bipolar disorder (BIP: (Stahl et al., 2019)), major depressive disorder (MDD: (Wray et al., 2018)), posttraumatic stress disorder (PTSD: (Nievergelt et al., 2019)); and for alcohol and tobacco use (Liu et al., 2019). For the marker gene sets, we used any genes defined as enriched per subpopulation (using marker test set *2*, from above), at the Benjamini & Hochberg false discovery rate (FDR) < 1e-12 (Benjamini and Hochberg, 1995). SNPs were first annotated to genes, using window sizes from −10kb to +35kb of each gene, with the 1000 Genomes EUR reference panel, and gene-level analyses were performed, using provided summary statistics from each of the above listed GWAS (via https://www.med.unc.edu/pgc/download-results/) and the `snp-wise=mean` model, to test whether there was enrichment of genetic risk for disease/phenotype in each gene. Following this, we performed the default `competitive` gene set analysis with the 68 regionally-defined marker sets, testing for association of gene-level risk and whether genes were enriched/specific to each subpopulation. From the empirical *p*-value of the gene set analysis, we performed multiple test correction with both false-discovery rate (FDR) and the stricter Bonferroni procedure (threshold *p* < 6.8e-5) across all 680 (68 regionally-defined subpopulations and 10 GWAS phenotypes tested) tests. All genetic association test results are provided in **Table S6**.

### RNAscope single molecule fluorescent in situ hybridization (smFISH)

Fresh frozen NAc from two independent donors was sectioned at 10μm and stored at −80°C. *In situ* hybridization assays were performed with RNAscope technology utilizing the RNAscope Fluorescent Multiplex Kit V2 and 4-plex Ancillary Kit (Cat # 323100, 323120 ACD, Hayward, California) according to the manufacturer’s instructions. Briefly, tissue sections were fixed with a 10% neutral buffered formalin solution (Cat # HT501128 Sigma-Aldrich, St. Louis, Missouri) for 30 minutes at room temperature (RT), series dehydrated in ethanol, pretreated with hydrogen peroxide for 10 minutes at RT, and treated with protease IV for 30 minutes. Sections were incubated with 5 different probe combinations to assess MSN and inhibitory neuron subtypes: 1) “Square”: *DRD1, TAC1, RXFP2, GABRQ* (Cat 524991-C4, 310711-C3, 452201, 483171-C2, ACD, Hayward, California); 2) “Circle”: *DRD1, TAC1, CRHR2, RXFP1* (Cat 524991-C4, 310711-C3, 469621, 422821-C2); 3) “Triangle”: *DRD1, DRD2, TAC1, PENK* (Cat 524991-C4, 553991, 310711-C2, 548301-C3); 4) “Star”: *DRD1, DRD2, CRHR2, HTR7* (Cat 524991-C4, 553991-C3, 469621, 413041-C2). 5) “Swirl”: *PVALB, GAD1, PTHLH, KIT* (Cat 422181-C4, 404031-C3, PTHLH, 606401-C2). Following probe labeling, sections were stored overnight in 4x SSC (saline-sodium citrate) buffer. After amplification steps (AMP1-3), probes were fluorescently labeled with Opal Dyes (Perkin Elmer, Waltham, MA; 1:500) and stained with DAPI (4’,6-diamidino-2-phenylindole) to label the nucleus. Lambda stacks were acquired in *z*-series using a Zeiss LSM780 confocal microscope equipped with a 63x x 1.4NA objective, a GaAsP spectral detector, and 405, 488, 555, and 647 lasers as previously described (Maynard et al., 2020b). All lambda stacks were acquired with the same imaging settings and laser power intensities. For each subject, high magnification 63x images were randomly acquired in the NAc (n= 2 subjects, n=2 sections per subject, n=12 images per section). Following image acquisition, lambda stacks in *z*-series were linearly unmixed in Zen software (weighted; no autoscale) using reference emission spectral profiles previously created in Zen (Maynard et al., 2020b) and saved as Carl Zeiss Image “.*czi*” files. Images were segmented and quantitatively analyzed in MATLAB using *dotdotdot* software (Maynard et al., 2020b) and statistical analyses were performed in R v3.6.3.

For each of the five experiments, we combined ROI-level data from all respective images, and used data-driven cutoffs based on distributional overlap to determine binary expression levels (i.e. expressed or unexpressed) for each gene/channel. For each experiment, we calculated the Euclidean distance of the vector of expressed targeted genes in each ROI to the cell type-specific targeted designs, where distance = 0 meant that ROI matched a predicted snRNA-seq cell type cluster (shown as larger points in **Figures E** and **S1, S2, S3**, & **S5**).

- Circle: 1033 ROIs were quantified across 48 images taken from 4 tissue sections across from 2 donors (two sections/donor). 251 ROIs were classified as *DRD1*+ with >3 dots post lipofuscin masking, and among these ROIs, *RXFP1* and *CRHR2* expression was classified as >3 dots and *TAC1* expression was classified as >6 dots.
- Square: 1126 ROIs were quantified across 48 images taken from 4 tissue sections across from 2 donors (two sections/donor). 341 ROIs were classified as *DRD1*+ with >3 dots post lipofuscin masking, and among these ROIs, *RXFP2, GABRQ*, and *TAC1* expression was classified as >6 dots.
- Triangle: 1039 ROIs were quantified across 47 images taken from 4 tissue sections across from 2 donors (two sections/donor). 271 ROIs were classified as either *DRD1*+ or *DRD2*+ with >3 dots post lipofuscin masking in either gene, and among these ROIs, *TAC1* and *PENK* expression was classified as >6 dots.
- Star: 1003 ROIs were quantified across 44 images taken from 4 tissue sections across from 2 donors (two sections/donor). 482 ROIs were classified as either *DRD1*+ or *DRD2*+ with >3 dots (post lipofuscin masking) in either gene, and among these ROIs, *HTR7* and *CRHR21* expression was classified as >6 dots.
- Swirl: 989 ROIs were quantified across 44 images taken from 4 tissue sections across from 2 donors (two sections/donor). 212 ROIs were classified as *GAD1*+ inhibitory neurons with >6 dots post lipofuscin masking, and among these ROIs, *PVALB, KIT* and *PTHLH* expression was classified as >6 dots.

### Data and Software Availability

All the code for processing and analyzing the data is available on the GitHub repository https://github.com/LieberInstitute/10xPilot_snRNAseq-human. The processed results files are hosted on Amazon S3 and the links are available on the README.md in the GitHub repository. For each of the five brain regions, we created an interactive website with the data using *iSEE* (*Rue-Albrecht et al., 2018*) and deployed at the LIBD shinyapps.io account at URLs such as https://libd.shinyapps.io/tran2020_Amyg/ (tran2020_sACC, tran2020_DLPFC, tran2020_NAc, tran2020_HPC).

## Acknowledgements

The authors would like to express their gratitude to our colleagues whose efforts have led to the donation of postmortem tissue to advance these studies, including at the Office of the Chief Medical Examiner of the State of Maryland, Baltimore Maryland. We also would like to acknowledge the contributions of Llewellyn B. Bigelow, MD and Amy Deep-Soboslay for their diagnostic expertise, and Daniel R. Weinberger for providing constructive commentary and editing of the manuscript. Finally, we are indebted to the generosity of the families of the decedents, who donated the brain tissue used in these studies. We thank the Johns Hopkins University Sidney Kimmel Comprehensive Cancer Center (SKCCC) Flow Cytometry Core, especially Jessica Gucwa, and the Johns Hopkins University Transcriptomics and Deep Sequencing Core, especially Linda Orzolek and Haiping Hao, for supporting snRNA-seq data generation. We also thank members of the laboratories of Dr. Weizhe Hong (UCLA) and Dr. Jeremy Day (University of Alabama) for facilitating data sharing and helpful conversations about cross-species comparisons.

## Funding

BKB, DBH, KM and AEJ were partially supported by R01DA042090; KRM, AS, LC-T, KM and AEJ were partially supported by 1R01MH123183. SCH was partially supported by R00HG009007. This project was also supported by the Lieber Institute for Brain Development.

## Author contribution

- M.N.T. - Conceptualization, Methodology, Data Curation, Investigation, Formal analysis, Visualization, Writing
- K.R.M. - Conceptualization, Methodology, Data Curation, Validation, Investigation, Writing, Visualization
- A.S. - Validation, Data Curation, Visualization
- L.C-T. - Methodology, Formal analysis, Data Curation, Writing, Visualization
- V.S. - Formal analysis, Visualization
- M.T. - Formal analysis, Visualization
- B.K.B. - Methodology, Visualization
- D.B.H. - Project administration, Funding acquisition
- S.C.H. - Methodology
- J.E.K. - Resources
- T.M.H. - Methodology, Resources
- K.M. - Conceptualization, Methodology, Writing, Supervision, Project administration, Funding acquisition
- A.E.J.- Conceptualization, Methodology, Software, Formal analysis, Writing, Visualization, Supervision, Project administration, Funding acquisition

## Supplementary Material

### Supplementary Tables

Table S1. Demographic/Sample Info

Table S2. Cell Ranger QC and Mapping Statistics

Table S3. Regionally-defined Nuclear Subpopulations, Donor-stratified

Table S4a-e. Top 40 Marker Lists (Both Pairwise and Cluster-vs-All-Others Results)

Table S5. Pan-brain Cluster Annotations, Region-stratified

Table S6. MAGMA association statistics across all regionally-defined subpopulations

### Supplementary Figures

**Figure S1.**
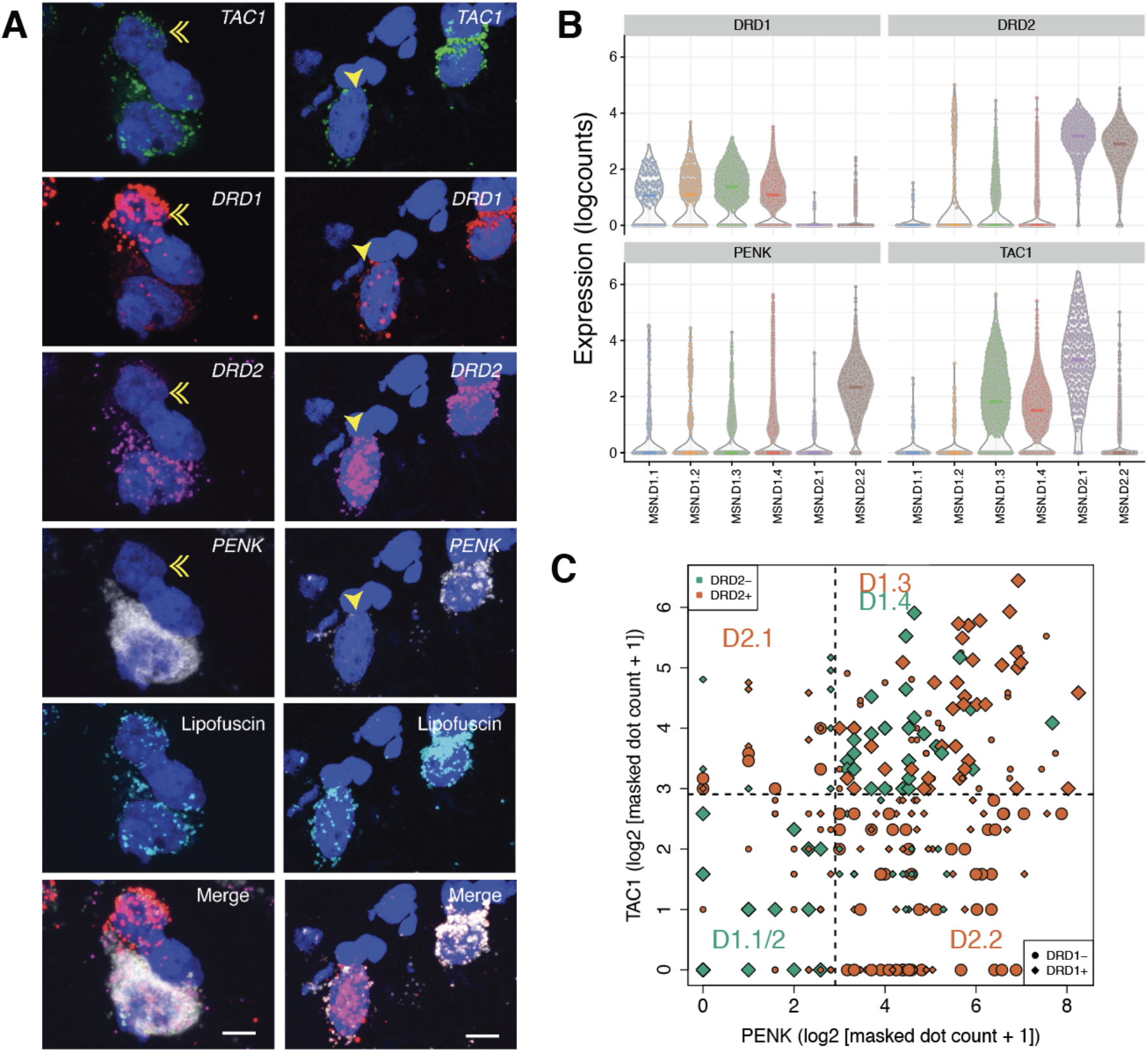
Differential expression of neuropeptide genes *TAC1* and *PENK* in D1 and D2 MSN subpopulations. Multiplex single molecule fluorescent in situ hybridization (smFISH) in human NAc. (**A**) Maximum intensity confocal projections showing expression of DAPI (nuclei), *DRD1, DRD2, TAC1, and PENK* and lipofuscin autofluorescence in two separate fields. Merged image without lipofuscin autofluorescence. Scale bar=10 μm. Double arrow indicates TAC1 negative D1 MSN. Single arrow indicates dual D1 and D2-expressing MSN. **(B)** Corresponding violin plots showing differential expression of *TAC1* and *PENK* in D1 and D2 MSNs subpopulations. **(C)** Log_2_ expression of respective transcript counts per smFISH ROI, post lipofuscin-masking (autofluorescence). Shape/color denote by *DRD1/DRD2* expression, respectively, and are enlarged where the Euclidean distance = 0 for prediction of MSN subclass for the respective ROI.

**Figure S2.**
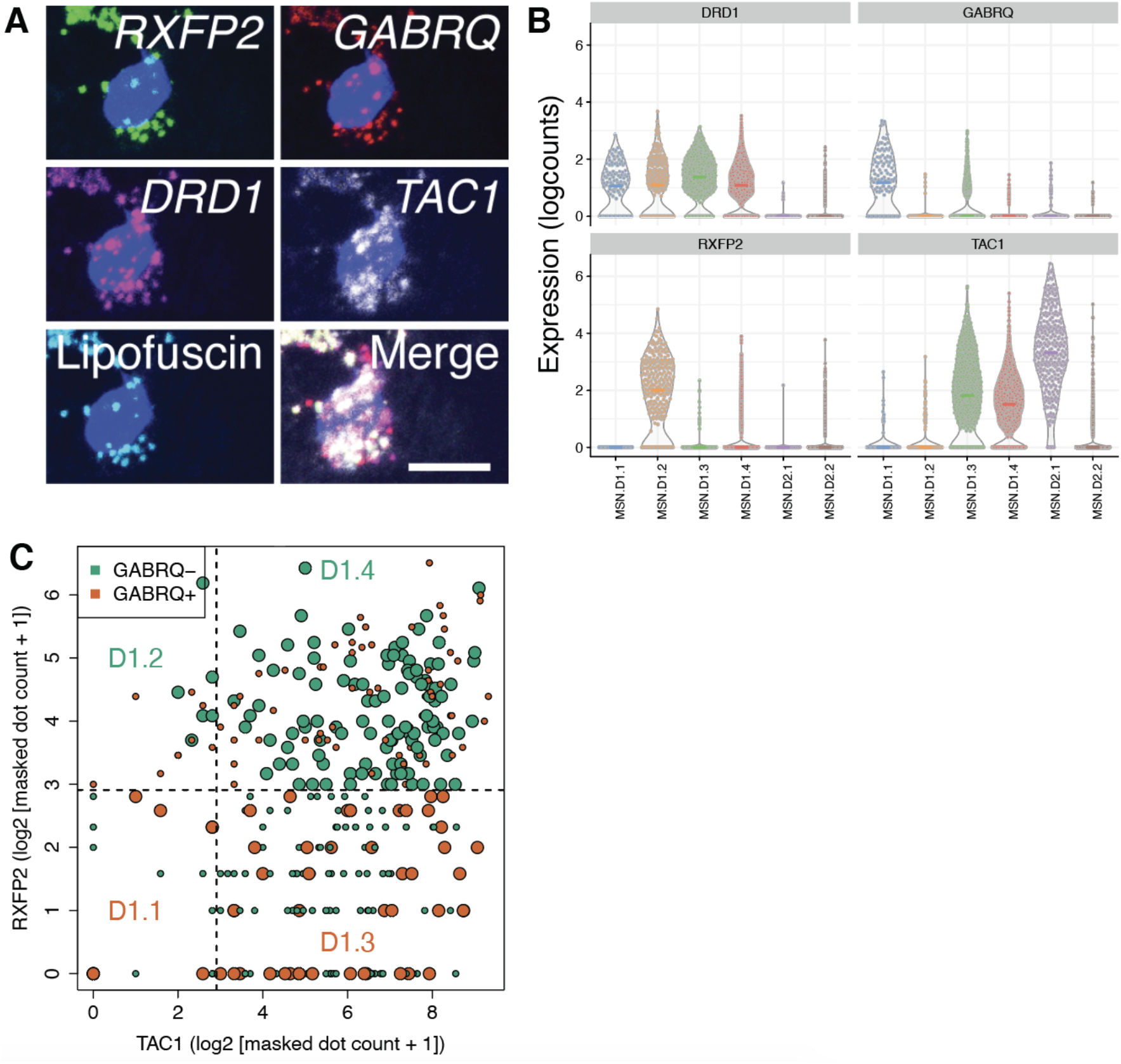
Further validation of D1 MSN subpopulations using smFISH. **(A)** Multiplex single molecule fluorescent in situ hybridization (smFISH) in human NAc depicting D1.3 MSN. Maximum intensity confocal projections showing expression of DAPI (nuclei), *RXFP2, GABRQ, DRD1, TAC1* and lipofuscin autofluorescence. Merged image without lipofuscin autofluorescence. Scale bar=10 μm. **(B)** Corresponding violin plots showing differential expression of these three genes in specific D1 subpopulations by snRNAseq. **(C)** Log_2_ expression of respective transcript counts per smFISH ROI, post lipofuscin-masking (autofluorescence). Points are colored by *GABRQ* expression and are enlarged where the Euclidean distance = 0 for prediction of MSN subclass for the respective ROI.

**Figure S3.**
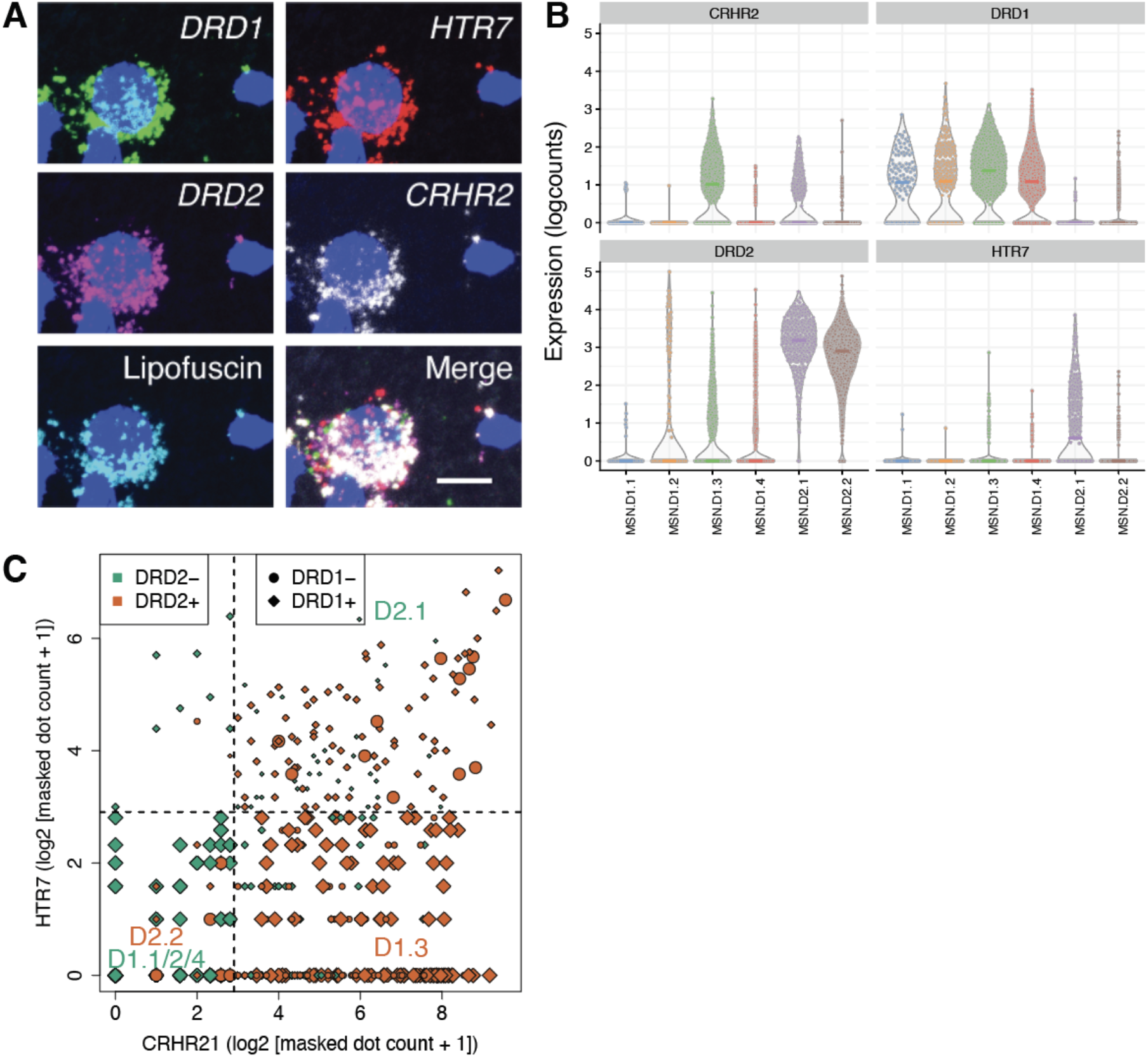
Confirmation of *HTR7*-expressing D2 MSNs in human NAc by smFISH. **(A)** Multiplex single molecule fluorescent in situ hybridization (smFISH) in human NAc depicting expression of *HTR7* in D2 MSNs. Maximum intensity confocal projections showing expression of DAPI (nuclei), *DRD1, HTR7, DRD2, CRHR2* and lipofuscin autofluorescence. Merged image without lipofuscin autofluorescence. Scale bar=10 μm. **(B)** Corresponding violin plots showing differential expression of *HTR7* and *CRHR2* in D1 and D2 MSNs subpopulations by snRNA-seq. **(C)** Log_2_ expression of respective transcript counts per smFISH ROI post lipofuscin-masking (autofluorescence). Shape/color denote by *DRD1/DRD2* expression, respectively, and are enlarged where the Euclidean distance = 0 for prediction of MSN subclass for the respective ROI.

**Figure S4.**
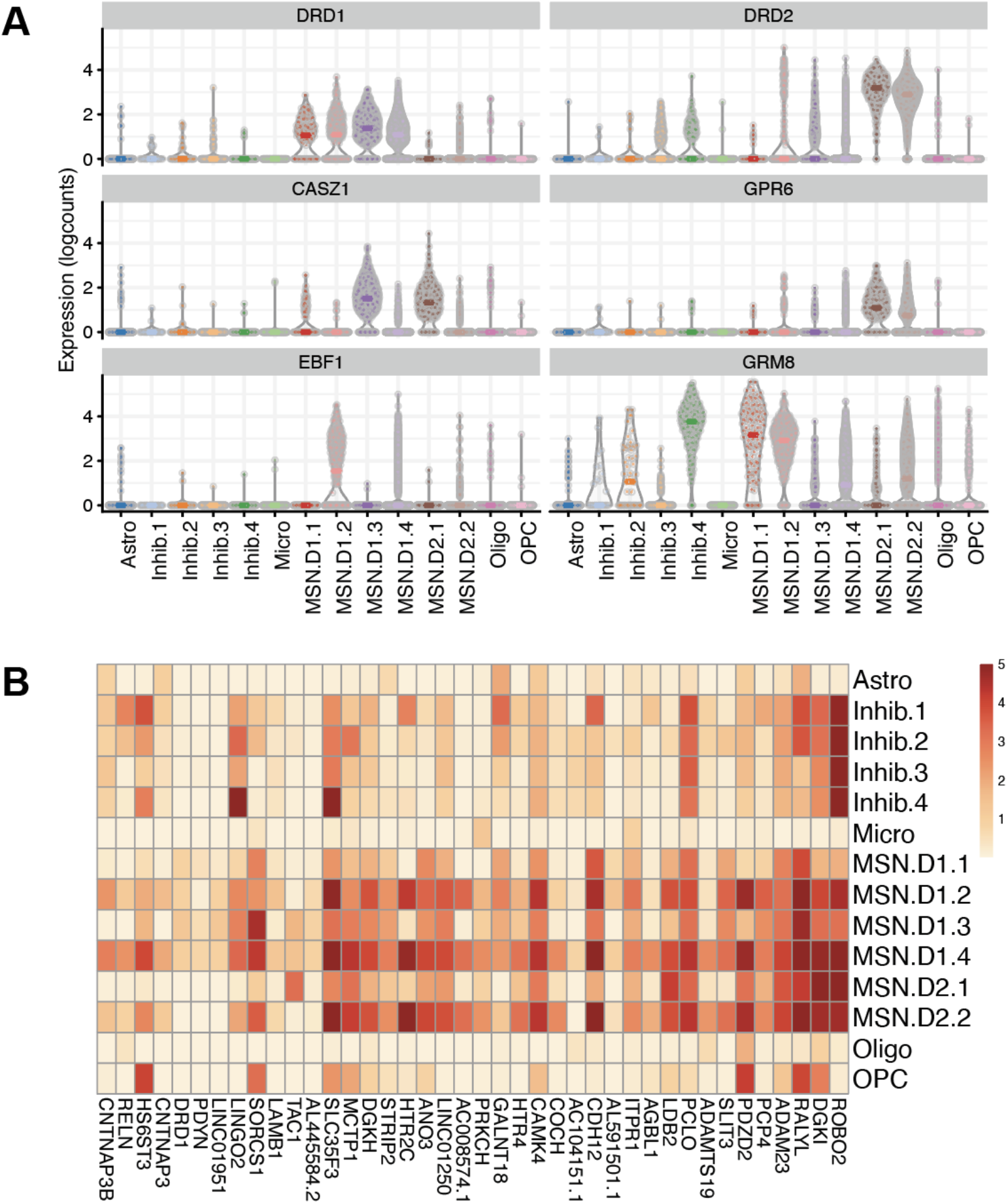
Other differentially expressed MSN markers and similarity between largest D1/D2 subpopulations. **(A)** Log_2_-normalized counts of other markers for MSN subpopulations not prioritized for smFISH validation, as above. GRM8 is included to show specific enrichment in D1.1 and D1.2. **(B)** Heatmap of mean snRNA-seq expression, showing top 40 markers for D1.4 and a similar pattern of expression in D2.2 subpopulation (scale thresholded to log_2_-normalized counts = 5.0).

**Figure S5.**
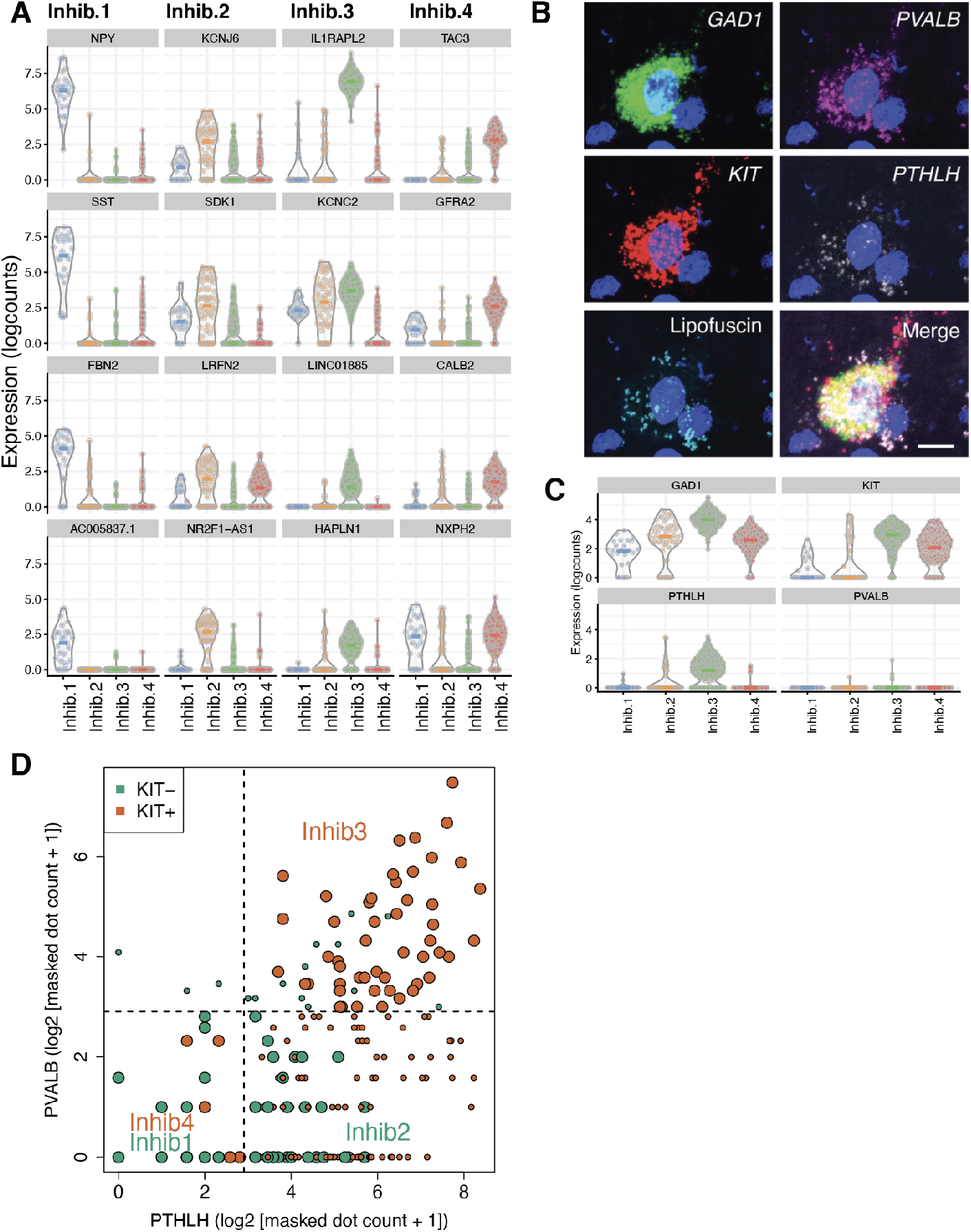
Characterization of interneuron subpopulations in human NAc. (**A)** Violin plots depicting top 4 genes (columns) in each interneuron subpopulation in NAc snRNA-seq. **(B)** Multiplex single molecule fluorescent in situ hybridization (smFISH) in human NAc depicting coexpression of *PVALB, KIT*, and *PTHLH* in *GAD1*+ interneurons. Maximum intensity confocal projections showing expression of DAPI (nuclei), *GAD1, PVALB, KIT, PTHLH* and lipofuscin autofluorescence. Merged image without lipofuscin autofluorescence. Scale bar=10 μm. (**C)** Corresponding violin plots showing expression of these genes in different interneuron subpopulations by snRNA-seq. **(D)** Log_2_ expression of respective transcript counts per smFISH ROI, post lipofuscin-masking (autofluorescence). Points are colored by *KIT* expression and are enlarged where the Euclidean distance = 0 for prediction of interneuron subclass for the respective ROI.

**Figure S6.**
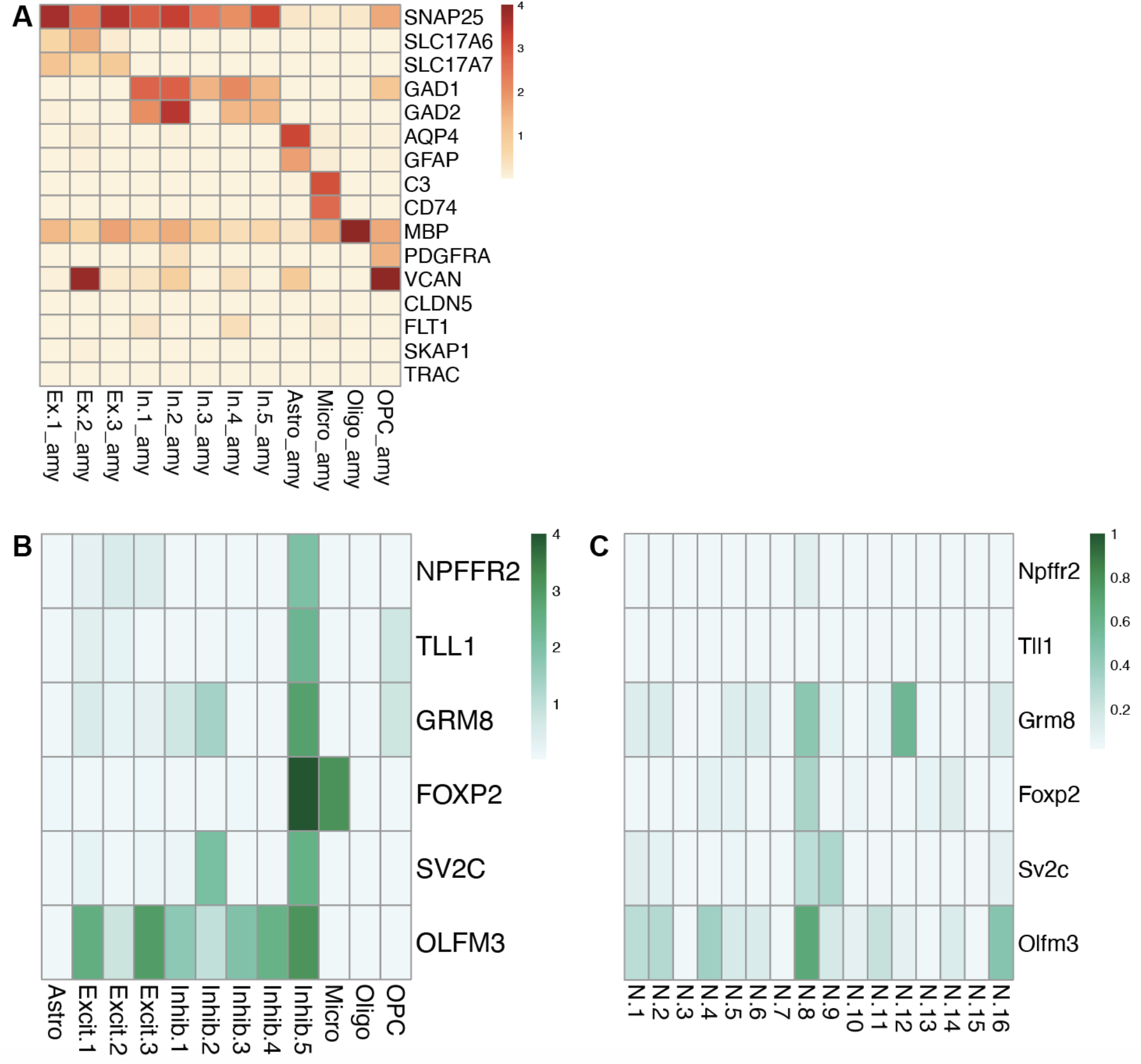
Broad cell type marker expression for AMY subpopulations and ‘Inhib.5’ vs. corresponding MeA ‘N.8’ shared markers. **(A)** Mean log_2_-normalized expression for broad cell type markers, used for annotation of AMY subpopulations. **(B)** Mean expression of top enriched markers for human AMY subpopulation Inhib.5 shared with **(C)** mouse MeA neuronal subclusters, as reported in (Chen et al., 2019). *Tll1*, however, was not defined as a marker of MeA ‘N.8’.

**Figure S7.**
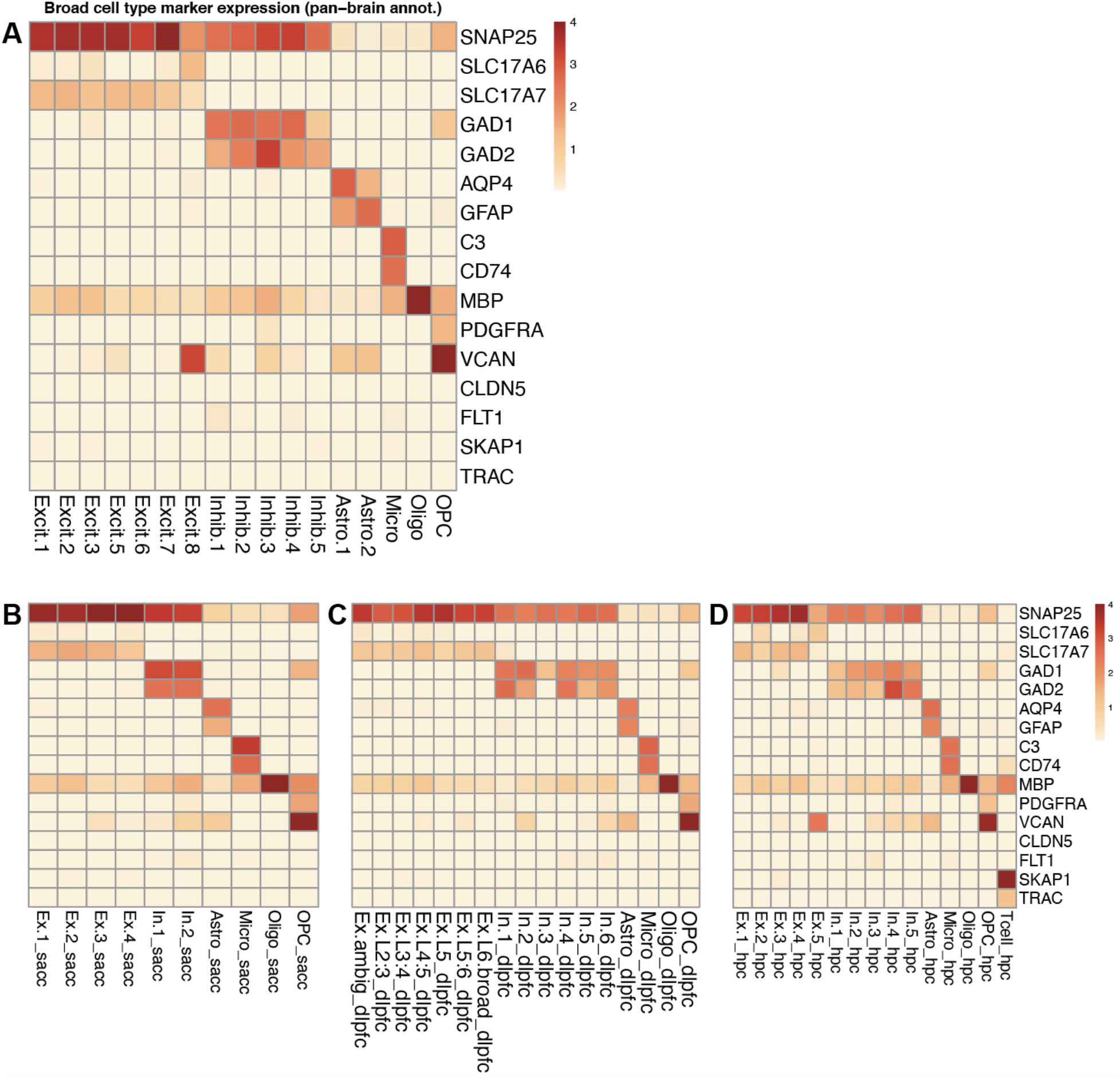
Broad cell type marker expression for pan-brain-defined clusters or regionally-defined populations. **(A)** Mean log_2_-normalized expression for broad cell type markers, used for annotation, in clusters defined across all nuclei in the five profiled regions. **(B)** Same as (A), but for regionally-defined subpopulations, for sACC, **(C)** DLPFC, and **(D)** HPC.

**Figure S8.**
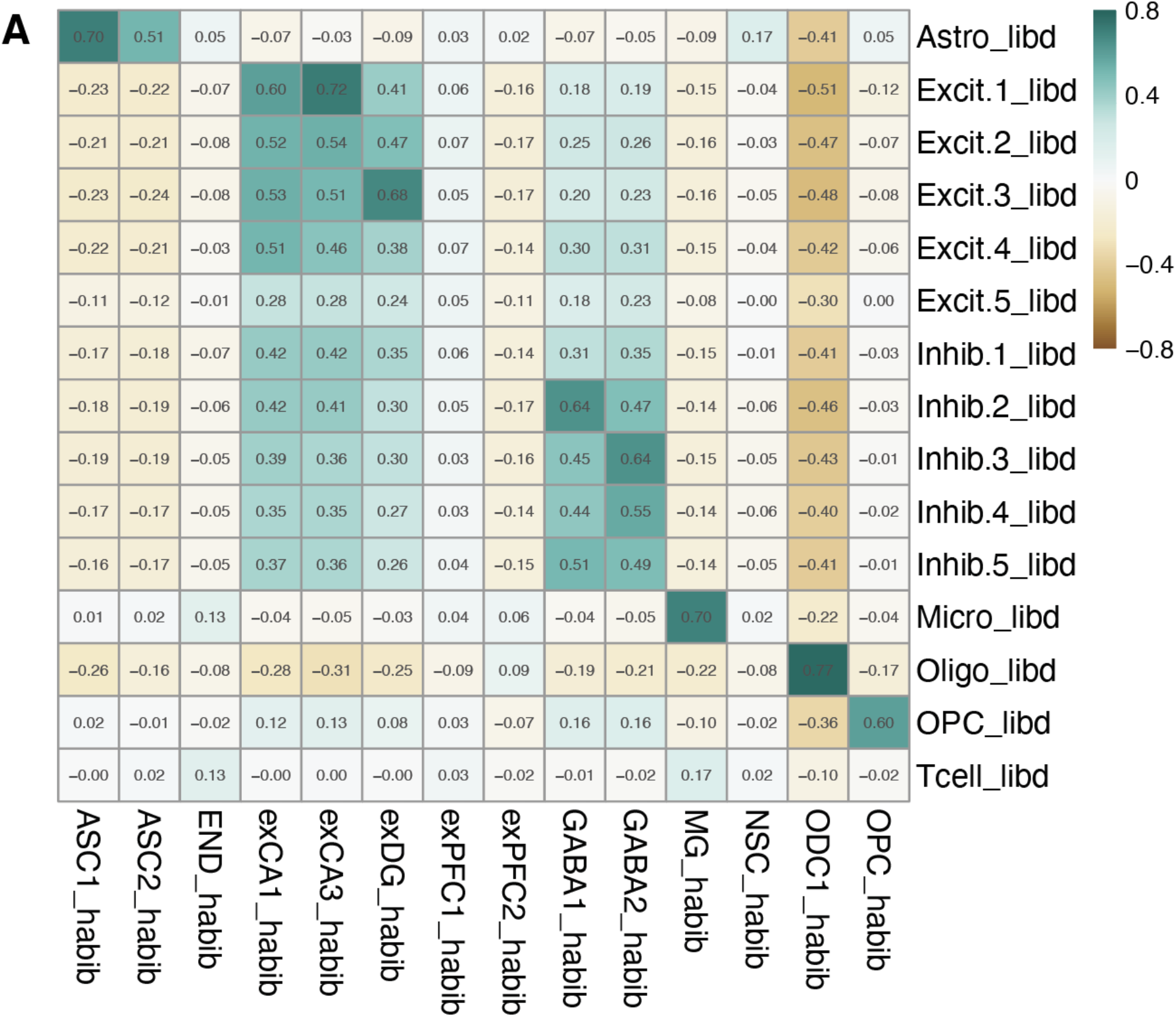
Benchmarking of HPC subpopulations to published data. **(A)** Correlation heatmap between HPC subclusters (rows) and the reported HPC populations in (Habib et al., 2017); columns). Printed values and scales show the Pearson correlation coefficient, correlating across all shared expressed genes and the *t*-statistics of their specificity test.

**Figure S9.**
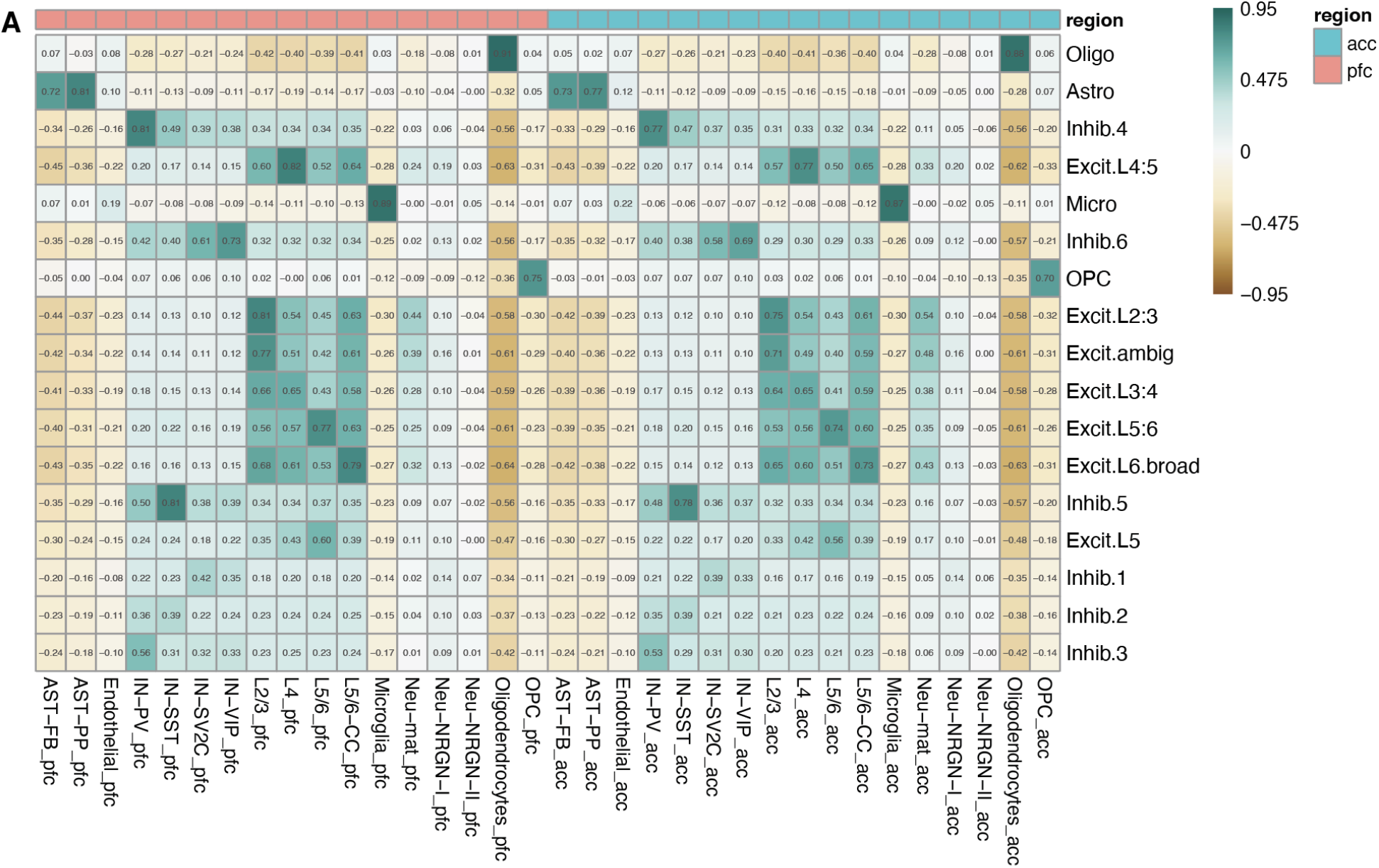
Benchmarking of DLPFC subpopulations to published data. **(A)** Correlation heatmap between DLPFC spatially-registered subpopulations (rows) and PFC 10x snRNA-seq data (columns) from (Velmeshev et al., 2019). Printed values and scales show the Pearson correlation coefficient, correlating across all shared expressed genes (26,970) and the *t*-statistics of their specificity test.

**Figure S10.**
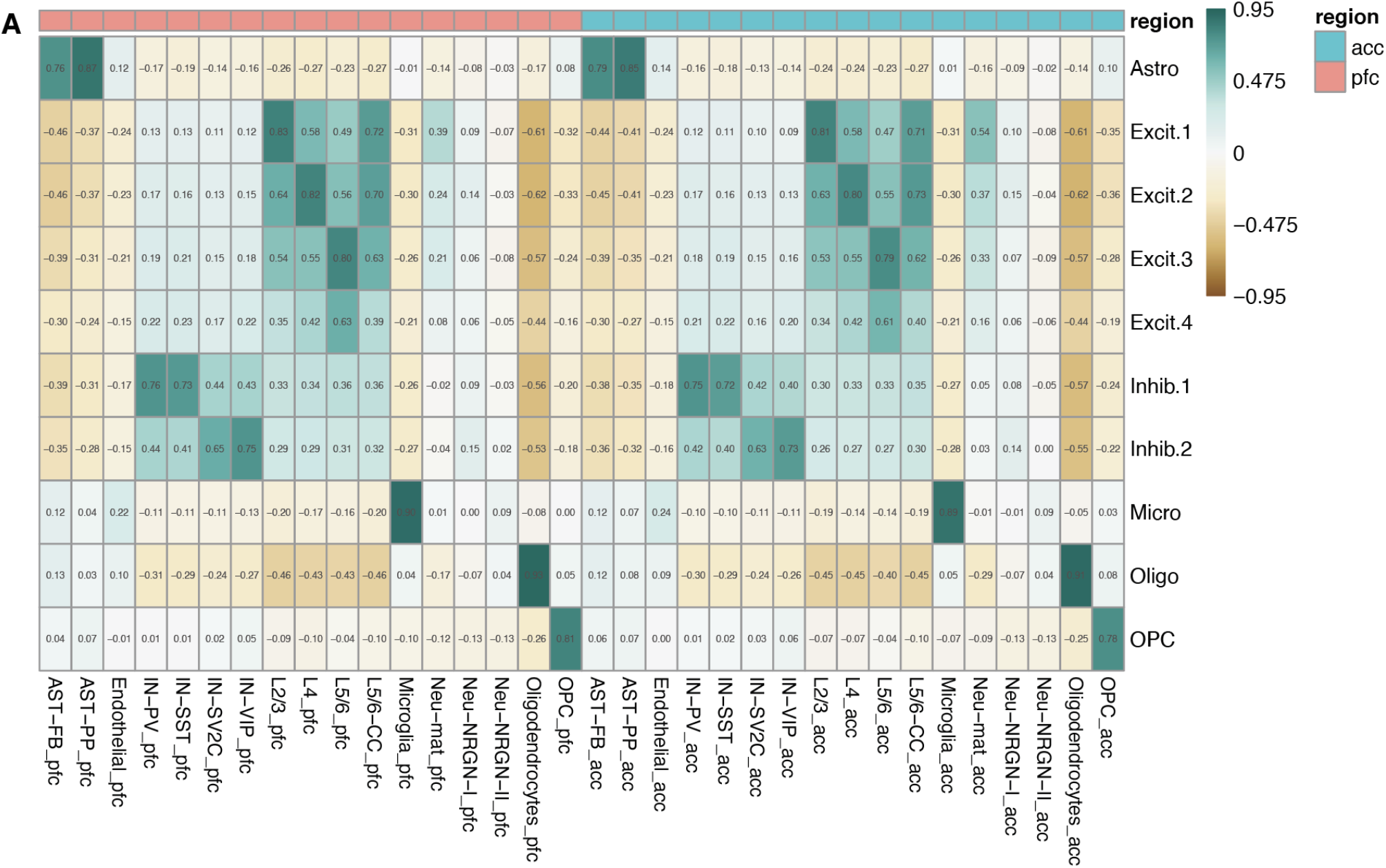
Benchmarking of sACC subpopulations to published data. **(A)** Correlation heatmap between sACC subpopulations (rows) and ACC 10x snRNA-seq data (columns) from (Velmeshev et al., 2019). Printed values and scales show the Pearson correlation coefficient, correlating across all shared expressed genes (27,422) and the *t*-statistics of their specificity test.

**Figure S11.**
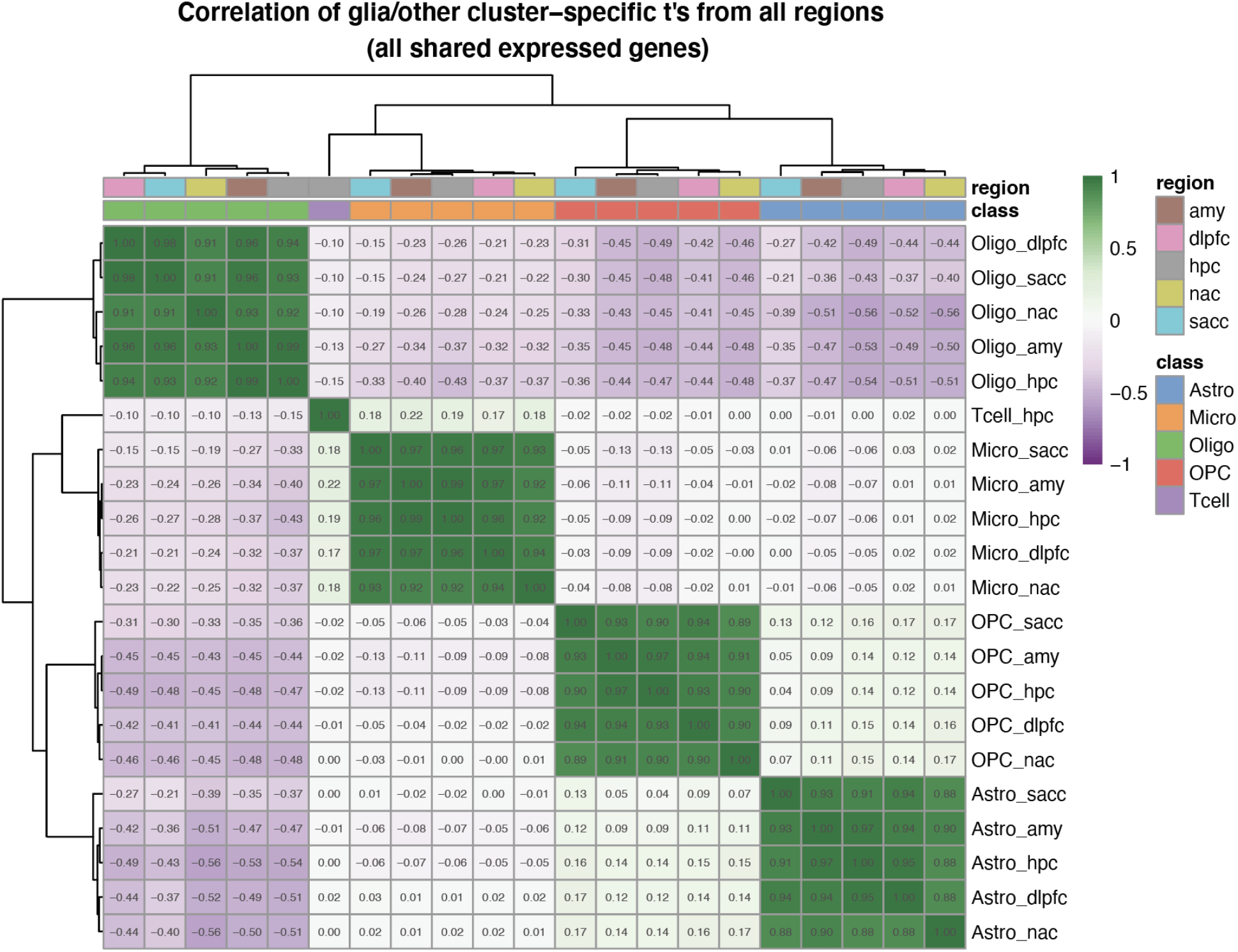
Comparison across all non-neuronal, regionally-defined subpopulations. Pairwise correlation of population-defined *t*-statistics, comparing 26,888 genes across the 21 non-neuronal subpopulations, collectively defined across each region (labeled in the suffix). Scale values are of Pearson correlation coefficient.

**Figure S12.**
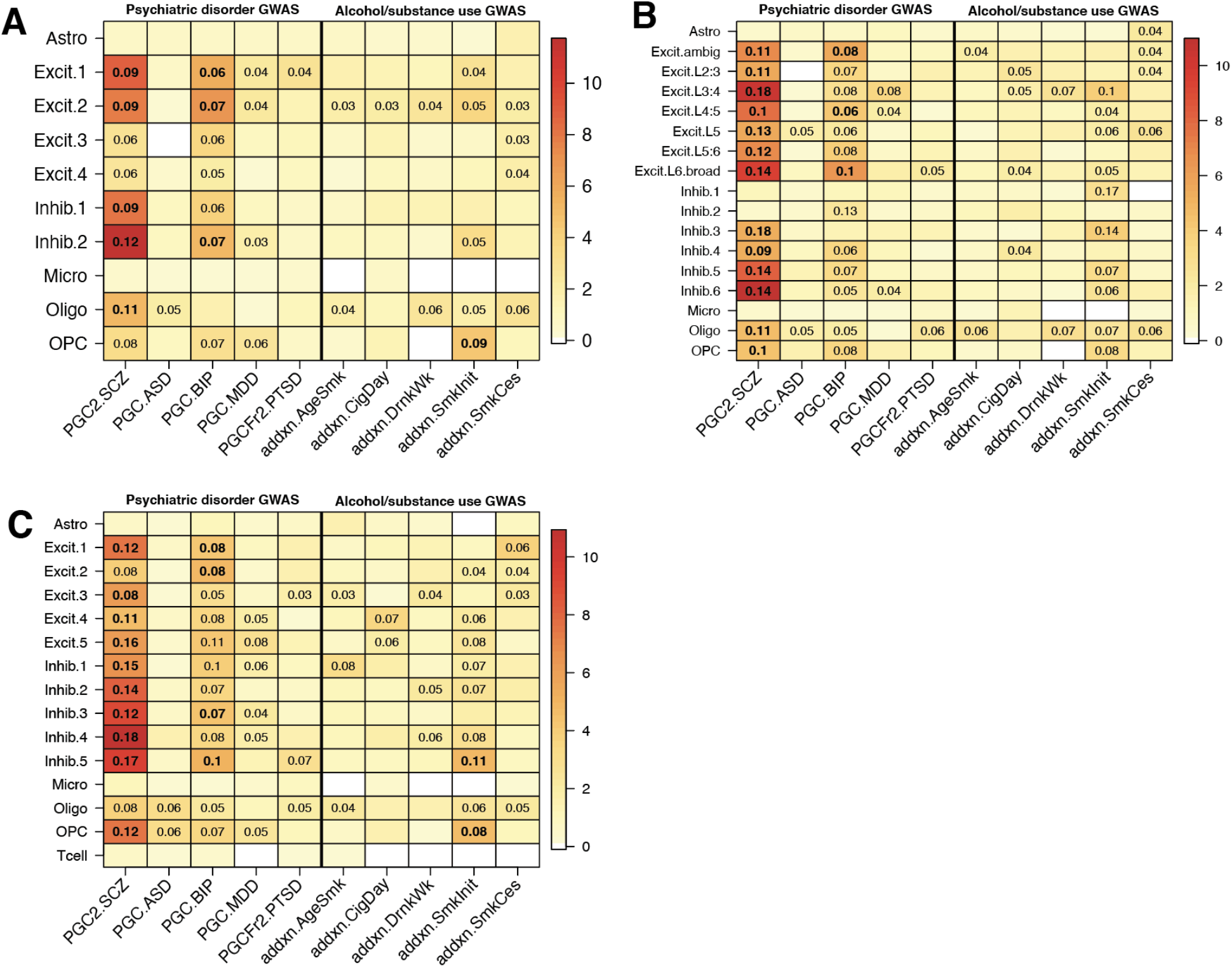
Genetic associations for HPC and cortical regions with psychiatric disease and addiction phenotypes. **(A)** MAGMA associations for each of 10 subpopulations profiled in sACC, **(B)** 17 spatially-resolved DLPFC subpopulations, and **(C)** 15 HPC subpopulations. Heatmap is colored by empirical −log10(*p-value*) for each association test. Displayed numbers are the effect size for significant associations (controlled for false discovery rate, FDR < 0.05), on a *Z* (standard normal) distribution. Bolded numbers are those that additionally satisfy a strict Bonferroni correction threshold of *p* < 6.8e-5.

## Notes

https://github.com/LieberInstitute/10xPilot_snRNAseq-human

### Competing Interest Statement

The authors have declared no competing interest.

## Bibliography

Amezquita, R.A., Lun, A.T.L., Becht, E., Carey, V.J., Carpp, L.N., Geistlinger, L., Marini, F., Rue-Albrecht, K., Risso, D., Soneson, C., et al. (2020). Orchestrating single-cell analysis with Bioconductor. Nat. Methods 17, 137–145.

Bailey, A.S., Willenbring, H., Jiang, S., Anderson, D.A., Schroeder, D.A., Wong, M.H., Grompe, M., and Fleming, W.H. (2006). Myeloid lineage progenitors give rise to vascular endothelium. Proc Natl Acad Sci USA 103, 13156–13161.

Barger, N., Stefanacci, L., Schumann, C.M., Sherwood, C.C., Annese, J., Allman, J.M., Buckwalter, J.A., Hof, P.R., and Semendeferi, K. (2012). Neuronal populations in the basolateral nuclei of the amygdala are differentially increased in humans compared with apes: a stereological study. J. Comp. Neurol. 520, 3035–3054.

Benjamini, Y., and Hochberg, Y. (1995). Controlling the False Discovery Rate: A Practical and Powerful Approach to Multiple Testing. J. R. Statist. Soc. B 57, 289–300.

Bonfiglio, J.J., Inda, C., Refojo, D., Holsboer, F., Arzt, E., and Silberstein, S. (2011). The corticotropin-releasing hormone network and the hypothalamic-pituitary-adrenal axis: molecular and cellular mechanisms involved. Neuroendocrinology 94, 12–20.

Bryois, J., Skene, N.G., Hansen, T.F., Kogelman, L.J.A., Watson, H.J., Liu, Z., Eating Disorders Working Group of the Psychiatric Genomics Consortium, International Headache Genetics Consortium, 23andMe Research Team, Brueggeman, L., et al. (2020). Genetic identification of cell types underlying brain complex traits yields insights into the etiology of Parkinson’s disease. Nat. Genet. 52, 482–493.

Chareyron, L.J., Banta Lavenex, P., Amaral, D.G., and Lavenex, P. (2011). Stereological analysis of the rat and monkey amygdala. J. Comp. Neurol. 519, 3218–3239.

Chen, P.B., Hu, R.K., Wu, Y.E., Pan, L., Huang, S., Micevych, P.E., and Hong, W. (2019). Sexually dimorphic control of parenting behavior by the medial amygdala. Cell 176, 1206–1221.e18.

Claes, S.J. (2004). Corticotropin-releasing hormone (CRH) in psychiatry: from stress to psychopathology. Ann. Med. 36, 50–61.

Collado-Torres, L., Burke, E.E., Peterson, A., Shin, J., Straub, R.E., Rajpurohit, A., Semick, S.A., Ulrich, W.S., BrainSeq Consortium, Price, A.J., et al. (2019). Regional Heterogeneity in Gene Expression, Regulation, and Coherence in the Frontal Cortex and Hippocampus across Development and Schizophrenia. Neuron 103, 203–216.e8.

Crowell, H.L., Soneson, C., Germain, P.-L., Calini, D., Collin, L., Raposo, C., Malhotra, D., and Robinson, M.D. (2019). On the discovery of population-specific state transitions from multisample multi-condition single-cell RNA sequencing data. BioRxiv.

Csardi, G., and Nepusz, T. (2006). The igraph software package for complex network research. InterJournal Complex Systems, 1695.

Darmanis, S., Sloan, S.A., Zhang, Y., Enge, M., Caneda, C., Shuer, L.M., Hayden Gephart, M.G., Barres, B.A., and Quake, S.R. (2015). A survey of human brain transcriptome diversity at the single cell level. Proc Natl Acad Sci USA 112, 7285–7290.

Enterría-Morales, D., Del Rey, N.L.-G., Blesa, J., López-López, I., Gallet, S., Prévot, V., López-Barneo, J., and d’Anglemont de Tassigny, X. (2020). Molecular targets for endogenous glial cell line-derived neurotrophic factor modulation in striatal parvalbumin interneurons. Brain Commun. 2, fcaa105.

Fenster, R.J., Lebois, L.A.M., Ressler, K.J., and Suh, J. (2018). Brain circuit dysfunction in post-traumatic stress disorder: from mouse to man. Nat. Rev. Neurosci. 19, 535–551.

Figueiro-Silva, J., Gruart, A., Clayton, K.B., Podlesniy, P., Abad, M.A., Gasull, X., Delgado-García, J.M., and Trullas, R. (2015). Neuronal pentraxin 1 negatively regulates excitatory synapse density and synaptic plasticity. J. Neurosci. 35, 5504–5521.

Finucane, H.K., Bulik-Sullivan, B., Gusev, A., Trynka, G., Reshef, Y., Loh, P.-R., Anttila, V., Xu, H., Zang, C., Farh, K., et al. (2015). Partitioning heritability by functional annotation using genome-wide association summary statistics. Nat. Genet. 47, 1228–1235.

Franjic, D., Choi, J., Skarica, M., Xu, C., Li, Q., Ma, S., Tebbenkamp, A.T.N., Santpere, G., Arellano, J.I., Gudelj, I., et al. (2020). Molecular diversity among adult hippocampal and entorhinal cells. BioRxiv.

Garrett, A., and Chang, K. (2008). The role of the amygdala in bipolar disorder development. Dev. Psychopathol. 20, 1285–1296.

Gerfen, C.R., Engber, T.M., Mahan, L.C., Susel, Z., Chase, T.N., Monsma, F.J., and Sibley, D.R. (1990). D1 and D2 dopamine receptor-regulated gene expression of striatonigral and striatopallidal neurons. Science 250, 1429–1432.

Gokce, O., Stanley, G.M., Treutlein, B., Neff, N.F., Camp, J.G., Malenka, R.C., Rothwell, P.E., Fuccillo, M.V., Südhof, T.C., and Quake, S.R. (2016). Cellular Taxonomy of the Mouse Striatum as Revealed by Single-Cell RNA-Seq. Cell Rep. 16, 1126–1137.

Graveland, G.A., Williams, R.S., and DiFiglia, M. (1985). A Golgi study of the human neostriatum: neurons and afferent fibers. J. Comp. Neurol. 234, 317–333.

Griffiths, J.A., Richard, A.C., Bach, K., Lun, A.T.L., and Marioni, J.C. (2018). Detection and removal of barcode swapping in single-cell RNA-seq data. Nat. Commun. 9, 2667.

Grove, J., Ripke, S., Als, T.D., Mattheisen, M., Walters, R.K., Won, H., Pallesen, J., Agerbo, E., Andreassen, O.A., Anney, R., et al. (2019). Identification of common genetic risk variants for autism spectrum disorder. Nat. Genet. 51, 431–444.

Grubman, A., Chew, G., Ouyang, J.F., Sun, G., Choo, X.Y., McLean, C., Simmons, R.K., Buckberry, S., Vargas-Landin, D.B., Poppe, D., et al. (2019). A single-cell atlas of entorhinal cortex from individuals with Alzheimer’s disease reveals cell-type-specific gene expression regulation. Nat. Neurosci. 22, 2087–2097.

Haber, S.N., and Knutson, B. (2010). The reward circuit: linking primate anatomy and human imaging. Neuropsychopharmacology 35, 4–26.

Habib, N., Li, Y., Heidenreich, M., Swiech, L., Avraham-Davidi, I., Trombetta, J.J., Hession, C., Zhang, F., and Regev, A. (2016). Div-Seq: Single-nucleus RNA-Seq reveals dynamics of rare adult newborn neurons. Science 353, 925–928.

Habib, N., Avraham-Davidi, I., Basu, A., Burks, T., Shekhar, K., Hofree, M., Choudhury, S.R., Aguet, F., Gelfand, E., Ardlie, K., et al. (2017). Massively parallel single-nucleus RNA-seq with DroNc-seq. Nat. Methods 14, 955–958.

Han, X., Zhou, Z., Fei, L., Sun, H., Wang, R., Chen, Y., Chen, H., Wang, J., Tang, H., Ge, W., et al. (2020). Construction of a human cell landscape at single-cell level. Nature 581, 303–309.

Heilig, M., and Koob, G.F. (2007). A key role for corticotropin-releasing factor in alcohol dependence. Trends Neurosci. 30, 399–406.

Heimer, L., Zahm, D.S., Churchill, L., Kalivas, P.W., and Wohltmann, C. (1991). Specificity in the projection patterns of accumbal core and shell in the rat. Neuroscience 41, 89–125.

Hodge, R.D., Bakken, T.E., Miller, J.A., Smith, K.A., Barkan, E.R., Graybuck, L.T., Close, J.L., Long, B., Johansen, N., Penn, O., et al. (2019). Conserved cell types with divergent features in human versus mouse cortex. Nature 573, 61–68.

Horii-Hayashi, N., Okuda, H., Tatsumi, K., Ishizaka, S., Yoshikawa, M., and Wanaka, A. (2008). Localization of chondroitin sulfate proteoglycan versican in adult brain with special reference to large projection neurons. Cell Tissue Res. 334, 163–177.

Howard, D.M., Adams, M.J., Clarke, T.-K., Hafferty, J.D., Gibson, J., Shirali, M., Coleman, J.R.I., Hagenaars, S.P., Ward, J., Wigmore, E.M., et al. (2019). Genome-wide meta-analysis of depression identifies 102 independent variants and highlights the importance of the prefrontal brain regions. Nat. Neurosci. 22, 343–352.

Huber, W., Carey, V.J., Gentleman, R., Anders, S., Carlson, M., Carvalho, B.S., Bravo, H.C., Davis, S., Gatto, L., Girke, T., et al. (2015). Orchestrating high-throughput genomic analysis with Bioconductor. Nat. Methods 12, 115–121.

Hu, P., Fabyanic, E., Kwon, D.Y., Tang, S., Zhou, Z., and Wu, H. (2017). Dissecting Cell-Type Composition and Activity-Dependent Transcriptional State in Mammalian Brains by Massively Parallel Single-Nucleus RNA-Seq. Mol. Cell 68, 1006–1015.e7.

Jaffe, A.E., Hoeppner, D.J., Saito, T., Blanpain, L., Ukaigwe, J., Burke, E.E., Collado-Torres, L., Tao, R., Tajinda, K., Maynard, K.R., et al. (2020). Profiling gene expression in the human dentate gyrus granule cell layer reveals insights into schizophrenia and its genetic risk. Nat. Neurosci. 23, 510–519.

Janak, P.H., and Tye, K.M. (2015). From circuits to behaviour in the amygdala. Nature 517, 284–292.

Kang, H.M., Subramaniam, M., Targ, S., Nguyen, M., Maliskova, L., McCarthy, E., Wan, E., Wong, S., Byrnes, L., Lanata, C.M., et al. (2018). Multiplexed droplet single-cell RNA-sequencing using natural genetic variation. Nat. Biotechnol. 36, 89–94.

Kawaguchi, Y. (1997). Neostriatal cell subtypes and their functional roles. Neurosci. Res. 27, 1–8.

Kronman, H., Richter, F., Labonté, B., Chandra, R., Zhao, S., Hoffman, G., Lobo, M.K., Schadt, E.E., and Nestler, E.J. (2019). Biology and Bias in Cell Type-Specific RNAseq of Nucleus Accumbens Medium Spiny Neurons. Sci. Rep. 9, 8350.

Lacar, B., Linker, S.B., Jaeger, B.N., Krishnaswami, S.R., Barron, J.J., Kelder, M.J.E., Parylak, S.L., Paquola, A.C.M., Venepally, P., Novotny, M., et al. (2016). Nuclear RNA-seq of single neurons reveals molecular signatures of activation. Nat. Commun. 7, 11022.

Lake, B.B., Ai, R., Kaeser, G.E., Salathia, N.S., Yung, Y.C., Liu, R., Wildberg, A., Gao, D., Fung, H.-L., Chen, S., et al. (2016). Neuronal subtypes and diversity revealed by single-nucleus RNA sequencing of the human brain. Science 352, 1586–1590.

Langfelder, P., Zhang, B., and with contributions from Steve Horvath (2016). dynamicTreeCut: Methods for Detection of Clusters in Hierarchical Clustering Dendrograms.

de Leeuw, C.A., Mooij, J.M., Heskes, T., and Posthuma, D. (2015). MAGMA: generalized geneset analysis of GWAS data. PLoS Comput. Biol. 11, e1004219.

Lin, Y.-T., Liu, T.-Y., Yang, C.-Y., Yu, Y.-L., Chen, T.-C., Day, Y.-J., Chang, C.-C., Huang, G.-J., and Chen, J.-C. (2016). Chronic activation of NPFFR2 stimulates the stress-related depressive behaviors through HPA axis modulation. Psychoneuroendocrinology 71, 73–85.

Lin, Y.-T., Yu, Y.-L., Hong, W.-C., Yeh, T.-S., Chen, T.-C., and Chen, J.-C. (2017). NPFFR2 activates the HPA axis and induces anxiogenic effects in rodents. Int. J. Mol. Sci. 18.

Lipska, B.K., Deep-Soboslay, A., Weickert, C.S., Hyde, T.M., Martin, C.E., Herman, M.M., and Kleinman, J.E. (2006). Critical factors in gene expression in postmortem human brain: Focus on studies in schizophrenia. Biol. Psychiatry 60, 650–658.

Liu, M., Jiang, Y., Wedow, R., Li, Y., Brazel, D.M., Chen, F., Datta, G., Davila-Velderrain, J., McGuire, D., Tian, C., et al. (2019). Association studies of up to 1.2 million individuals yield new insights into the genetic etiology of tobacco and alcohol use. Nat. Genet. 51, 237–244.

Li, M., Santpere, G., Imamura Kawasawa, Y., Evgrafov, O.V., Gulden, F.O., Pochareddy, S., Sunkin, S.M., Li, Z., Shin, Y., Zhu, Y., et al. (2018a). Integrative functional genomic analysis of human brain development and neuropsychiatric risks. Science 362.

Li, Z., Chen, Z., Fan, G., Li, A., Yuan, J., and Xu, T. (2018b). Cell-Type-Specific Afferent Innervation of the Nucleus Accumbens Core and Shell. Front. Neuroanat. 12, 84.

Lobo, M.K. (2009). Molecular profiling of striatonigral and striatopallidal medium spiny neurons past, present, and future. Int. Rev. Neurobiol. 89, 1–35.

Lobo, M.K., Karsten, S.L., Gray, M., Geschwind, D.H., and Yang, X.W. (2006). FACS-array profiling of striatal projection neuron subtypes in juvenile and adult mouse brains. Nat. Neurosci. 9, 443–452.

Lun, A.T.L., and Marioni, J.C. (2017). Overcoming confounding plate effects in differential expression analyses of single-cell RNA-seq data. Biostatistics 18, 451–464.

Lun, A.T.L., McCarthy, D.J., and Marioni, J.C. (2016). A step-by-step workflow for low-level analysis of single-cell RNA-seq data with Bioconductor. [version 2; peer review: 3 approved, 2 approved with reservations]. F1000Res. 5, 2122.

Lun, A.T.L., Riesenfeld, S., Andrews, T., Dao, T.P., Gomes, T., participants in the 1st Human Cell Atlas Jamboree, and Marioni, J.C. (2019). EmptyDrops: distinguishing cells from empty droplets in droplet-based single-cell RNA sequencing data. Genome Biol. 20, 63.

van der Maaten, L., and Hinton, G. (2008). Visualizing Data using t-SNE. J Mach Learn Res 9, 2579–2605.

Mathys, H., Davila-Velderrain, J., Peng, Z., Gao, F., Mohammadi, S., Young, J.Z., Menon, M., He, L., Abdurrob, F., Jiang, X., et al. (2019). Single-cell transcriptomic analysis of Alzheimer’s disease. Nature 570, 332–337.

Maynard, K.E., Collado-Torres, L., Weber, L.M., Uytingco, C., Barry, B.K., Williams, S.R., Catallini, J.L., Tran, M.N., Besich, Z., Tippani, M., et al. (2020a). Transcriptome-scale spatial gene expression in the human dorsolateral prefrontal cortex. BioRxiv.

Maynard, K.R., Tippani, M., Takahashi, Y., Phan, B.N., Hyde, T.M., Jaffe, A.E., and Martinowich, K. (2020b). dotdotdot: an automated approach to quantify multiplex single molecule fluorescent in situ hybridization (smFISH) images in complex tissues. Nucleic Acids Res.

McCarthy, D.J., Campbell, K.R., Lun, A.T.L., and Wills, Q.F. (2017). Scater: pre-processing, quality control, normalization and visualization of single-cell RNA-seq data in R. Bioinformatics 33, 1179–1186.

McInnes, L., Healy, J., and Melville, J. (2018). UMAP: Uniform Manifold Approximation and Projection. ArXiv.

Murray, E.A., Wise, S.P., and Drevets, W.C. (2011). Localization of dysfunction in major depressive disorder: prefrontal cortex and amygdala. Biol. Psychiatry 69, e43–54.

Nagy, C., Maitra, M., Tanti, A., Suderman, M., Théroux, J.-F., Davoli, M.A., Perlman, K., Yerko, V., Wang, Y.C., Tripathy, S.J., et al. (2020). Single-nucleus transcriptomics of the prefrontal cortex in major depressive disorder implicates oligodendrocyte precursor cells and excitatory neurons. Nat. Neurosci. 23, 771–781.

Nievergelt, C.M., Maihofer, A.X., Klengel, T., Atkinson, E.G., Chen, C.-Y., Choi, K.W., Coleman, J.R.I., Dalvie, S., Duncan, L.E., Gelernter, J., et al. (2019). International meta-analysis of PTSD genome-wide association studies identifies sex-and ancestry-specific genetic risk loci. Nat. Commun. 10, 4558.

Pardiñas, A.F., Holmans, P., Pocklington, A.J., Escott-Price, V., Ripke, S., Carrera, N., Legge, S.E., Bishop, S., Cameron, D., Hamshere, M.L., et al. (2018). Common schizophrenia alleles are enriched in mutation-intolerant genes and in regions under strong background selection. Nat. Genet. 50, 381–389.

Payer, D., Williams, B., Mansouri, E., Stevanovski, S., Nakajima, S., Le Foll, B., Kish, S., Houle, S., Mizrahi, R., George, S.R., et al. (2017). Corticotropin-releasing hormone and dopamine release in healthy individuals. Psychoneuroendocrinology 76, 192–196.

Prensa, L., Richard, S., and Parent, A. (2003). Chemical anatomy of the human ventral striatum and adjacent basal forebrain structures. J. Comp. Neurol. 460, 345–367.

Rizzardi, L.F., Hickey, P.F., Rodriguez DiBlasi, V., Tryggvadóttir, R., Callahan, C.M., Idrizi, A., Hansen, K.D., and Feinberg, A.P. (2019). Neuronal brain-region-specific DNA methylation and chromatin accessibility are associated with neuropsychiatric trait heritability. Nat. Neurosci. 22, 307–316.

Rue-Albrecht, K., Marini, F., Soneson, C., and Lun, A.T.L. (2018). iSEE: Interactive SummarizedExperiment Explorer. [version 1; peer review: 3 approved]. F1000Res. 7, 741.

Russo, S.J., and Nestler, E.J. (2013). The brain reward circuitry in mood disorders. Nat. Rev. Neurosci. 14, 609–625.

Rymar, V.V., Sasseville, R., Luk, K.C., and Sadikot, A.F. (2004). Neurogenesis and stereological morphometry of calretinin-immunoreactive GABAergic interneurons of the neostriatum. J. Comp. Neurol. 469, 325–339.

Salgado, S., and Kaplitt, M.G. (2015). The nucleus accumbens: A comprehensive review. Stereotact. Funct. Neurosurg. 93, 75–93.

Saunders, A., Macosko, E.Z., Wysoker, A., Goldman, M., Krienen, F.M., de Rivera, H., Bien, E., Baum, M., Bortolin, L., Wang, S., et al. (2018). Molecular Diversity and Specializations among the Cells of the Adult Mouse Brain. Cell 174, 1015–1030.e16.

Savell, K.E., Tuscher, J.J., Zipperly, M.E., Duke, C.G., Phillips, R.A., Bauman, A.J., Thukral, S., Sultan, F.A., Goska, N.A., Ianov, L., et al. (2020). A dopamine-induced gene expression signature regulates neuronal function and cocaine response. Sci. Adv. 6, eaba4221.

Schirmer, L., Velmeshev, D., Holmqvist, S., Kaufmann, M., Werneburg, S., Jung, D., Vistnes, S., Stockley, J.H., Young, A., Steindel, M., et al. (2019). Neuronal vulnerability and multilineage diversity in multiple sclerosis. Nature 573, 75–82.

Schizophrenia Working Group of the Psychiatric Genomics Consortium (2014). Biological insights from 108 schizophrenia-associated genetic loci. Nature 511, 421–427.

Schumann, C.M., and Amaral, D.G. (2005). Stereological estimation of the number of neurons in the human amygdaloid complex. J. Comp. Neurol. 491, 320–329.

Sindreu, C., and Storm, D.R. (2011). Modulation of neuronal signal transduction and memory formation by synaptic zinc. Front. Behav. Neurosci. 5, 68.

Skene, N.G., Bryois, J., Bakken, T.E., Breen, G., Crowley, J.J., Gaspar, H.A., Giusti-Rodriguez, P., Hodge, R.D., Miller, J.A., Muñoz-Manchado, A.B., et al. (2018). Genetic identification of brain cell types underlying schizophrenia. Nat. Genet. 50, 825–833.

Sorvari, H., Soininen, H., Paljärvi, L., Karkola, K., and Pitkänen, A. (1995). Distribution of parvalbumin-immunoreactive cells and fibers in the human amygdaloid complex. J. Comp. Neurol. 360, 185–212.

Stahl, E.A., Breen, G., Forstner, A.J., McQuillin, A., Ripke, S., Trubetskoy, V., Mattheisen, M., Wang, Y., Coleman, J.R.I., Gaspar, H.A., et al. (2019). Genome-wide association study identifies 30 loci associated with bipolar disorder. Nat. Genet. 51, 793–803.

Stanley, G., Gokce, O., Malenka, R.C., Südhof, T.C., and Quake, S.R. (2020). Continuous and discrete neuron types of the adult murine striatum. Neuron 105, 688–699.e8.

Tamura, G., Olson, D., Miron, J., and Clark, T.G. (2005). Tolloid-like 1 is negatively regulated by stress and glucocorticoids. Brain Res. Mol. Brain Res. 142, 81–90.

Tepper, J.M., and Bolam, J.P. (2004). Functional diversity and specificity of neostriatal interneurons. Curr. Opin. Neurobiol. 14, 685–692.

Thrupp, N., Sala Frigerio, C., Wolfs, L., Skene, N.G., Fattorelli, N., Poovathingal, S., Fourne, Y., Matthews, P.M., Theys, T., Mancuso, R., et al. (2020). Single-Nucleus RNA-Seq Is Not Suitable for Detection of Microglial Activation Genes in Humans. Cell Rep. 32, 108189.

Tyszka, J.M., and Pauli, W.M. (2016). In vivo delineation of subdivisions of the human amygdaloid complex in a high-resolution group template. Hum. Brain Mapp. 37, 3979–3998.

Velmeshev, D., Schirmer, L., Jung, D., Haeussler, M., Perez, Y., Mayer, S., Bhaduri, A., Goyal, N., Rowitch, D.H., and Kriegstein, A.R. (2019). Single-cell genomics identifies cell type-specific molecular changes in autism. Science 364, 685–689.

Voorn, P., Gerfen, C.R., and Groenewegen, H.J. (1989). Compartmental organization of the ventral striatum of the rat: immunohistochemical distribution of enkephalin, substance P, dopamine, and calcium-binding protein. J. Comp. Neurol. 289, 189–201.

Wassum, K.M., and Izquierdo, A. (2015). The basolateral amygdala in reward learning and addiction. Neurosci. Biobehav. Rev. 57, 271–283.

Wray, N.R., Ripke, S., Mattheisen, M., Trzaskowski, M., Byrne, E.M., Abdellaoui, A., Adams, M.J., Agerbo, E., Air, T.M., Andlauer, T.M.F., et al. (2018). Genome-wide association analyses identify 44 risk variants and refine the genetic architecture of major depression. Nat. Genet. 50, 668–681.

Wu, Y.E., Pan, L., Zuo, Y., Li, X., and Hong, W. (2017). Detecting Activated Cell Populations Using Single-Cell RNA-Seq. Neuron 96, 313–329.e6.

Xie, P., Kranzler, H.R., Yang, C., Zhao, H., Farrer, L.A., and Gelernter, J. (2013). Genome-wide association study identifies new susceptibility loci for posttraumatic stress disorder. Biol. Psychiatry 74, 656–663.

Yao, J.-J., Zhao, Q.-R., Lu, J.-M., and Mei, Y.-A. (2018). Functions and the related signaling pathways of the neurotrophic factor neuritin. Acta Pharmacol. Sin. 39, 1414–1420.

Yong, W., Spence, J.P., Eskay, R., Fitz, S.D., Damadzic, R., Lai, D., Foroud, T., Carr, L.G., Shekhar, A., Chester, J.A., et al. (2014). Alcohol-preferring rats show decreased corticotropin-releasing hormone-2 receptor expression and differences in HPA activation compared to alcohol-nonpreferring rats. Alcohol. Clin. Exp. Res. 38, 1275–1283.

Zahm, D.S., and Heimer, L. (1993). Specificity in the efferent projections of the nucleus accumbens in the rat: comparison of the rostral pole projection patterns with those of the core and shell. J. Comp. Neurol. 327, 220–232.

Zeisel, A., Hochgerner, H., Lönnerberg, P., Johnsson, A., Memic, F., van der Zwan, J., Häring, M., Braun, E., Borm, L.E., La Manno, G., et al. (2018). Molecular architecture of the mouse nervous system. Cell 174, 999–1014.e22.

Zhang, X., Cheng, H., Zuo, Z., Zhou, K., Cong, F., Wang, B., Zhuo, Y., Chen, L., Xue, R., and Fan, Y. (2018). Individualized Functional Parcellation of the Human Amygdala Using a Semisupervised Clustering Method: A 7T Resting State fMRI Study. Front. Neurosci. 12, 270.

Zheng, G.X.Y., Terry, J.M., Belgrader, P., Ryvkin, P., Bent, Z.W., Wilson, R., Ziraldo, S.B., Wheeler, T.D., McDermott, G.P., Zhu, J., et al. (2017). Massively parallel digital transcriptional profiling of single cells. Nat. Commun. 8, 14049.

Zhong, S., Zhang, S., Fan, X., Wu, Q., Yan, L., Dong, J., Zhang, H., Li, L., Sun, L., Pan, N., et al. (2018). A single-cell RNA-seq survey of the developmental landscape of the human prefrontal cortex. Nature 555, 524–528.

Zhong, S., Ding, W., Sun, L., Lu, Y., Dong, H., Fan, X., Liu, Z., Chen, R., Zhang, S., Ma, Q., et al. (2020). Decoding the development of the human hippocampus. Nature 577, 531–536.

